# Realtime observation of ATP-driven single B_12_ molecule translocation through BtuCD-F

**DOI:** 10.1101/2022.12.02.518935

**Authors:** Lingwei Zhu, Jinrang Kim, Kun Leng, John E. Ramos, Colin D. Kinz-Thompson, Nathan K. Karpowich, Ruben L. Gonzalez, John F. Hunt

## Abstract

<u>A</u>TP-<u>B</u>inding <u>C</u>assette (ABC) Transporters use ATP binding and hydrolysis to power transmembrane transport of chemically diverse substrates. Current knowledge of their mechanism comes primarily from static structures of stable intermediates along the transport cycle. Recently, single-molecule fluorescence resonance energy transfer (smFRET) measurements have generated insight into the functional dynamics of transmembrane transporters, but studies to date lack direct information on the physical movement of the transport substrate. Here, we report development of an smFRET system that exploits fluorescence quenching by vitamin B_12_ to track its location in real time during ATP-driven transport by nanodisc-reconstituted *E. coli* BtuCD-F, an extensively studied type II ABC importer. Our data demonstrate that transmembrane translocation of B_12_ is driven by two sequential high-energy conformational changes that are inaccessible to standard structural methods because they are inherently transient. The first moves B_12_ from the periplasm into the transmembrane domain of the transporter; notably, this reaction is driven by hydrolysis of a single ATP molecule, in contrast to the mechanism established for several other ABC Transporter families in which ATP-binding drives the mechanochemical power-stroke prior to hydrolysis. The second mediates B_12_ release on the opposite side of the transporter, and it is driven by formation of a hyper-stable complex between BtuCD and BtuF. Hydrolysis of a second single ATP molecule is then required to dissociate BtuCD from the BtuF substrate-binding protein to enable it to bind B_12_ and initiate another round of transport. Our experiments have visualized substrate translocation in real-time at a single-molecule level and provided unprecedented information on the mechanism and dynamics of a paradigmatic transmembrane transport process.

---

The ABC protein superfamily is ubiquitous in all phylogenetic kingdoms and one of the largest families in the Protein Family (PFAM) database^**1**^. Most ABC proteins are transmembrane (TM) transporters that function as homodimeric or homologous heterodimeric complexes containing two transmembrane domains (TMDs) each coupled to a well-conserved ATP-binding cassette that is alternatively called a nucleotide-binding domain (NBD). Type I and II ABC importers also require a peripherally bound substrate-binding protein (SBP) that cycles on and off the periplasmic surface of the TMD to deliver the transport substrate. The binding and hydrolysis of two ATP molecules at the interface between the two NBDs drives the mechanochemistry of the transport process. Extensive studies demonstrate that ABC exporters and type I ABC importers both employ an alternating-site-access mechanism^**2**^ in which the binding site for the transport substrate is exposed to opposite sides of the membrane in the nucleotide-free *vs.* ATP bound conformations^3–10^. ATP binding drives conversion between these global conformations, and this mechanochemical “power-stroke” results in release of the substrate on the opposite side of the membrane from where it initially bound, while ATP hydrolysis gates relaxation back to the starting ground-state conformation to reset the transporter.

In contrast, the transport mechanism remains controversial for type II ABC importers^**11-15**^, which transport metal chelate complexes that are generally larger than the substrates transported by type I importers. The most extensively studied type II importer is the *E. coli* vitamin B_12_ transporter BtuCD-F^**12,13,15–26**^, in which BtuC is the TMD, BtuD is the NBD, and BtuF is the SBP (**Fig. 1c**). Crystallographic studies of BtuCD-F 18,24,25 show an internal cavity in its TMD that is slightly smaller than the B_12_ transport substrate (**Fig. S1a** in the ***Supplementary Information***). However, B_12_ has not been observed at this site in any structure, and there is no biochemical data establishing stable occupancy of this site by B_12_. Furthermore, X-ray crystallographic structures determined in both nucleotide-free and ATP-bound states show that this site is consistently sealed from both the cytoplasmic and periplasmic surfaces of the TMD (**Fig. S1a**), making it unclear how binding/hydrolysis of ATP drives transport. These observations, which contrast with extensive results on type I importers^**27-30**^, suggest that type II ABC importers like BtuCD-F employ a different transport mechanism that does not involve a mechanochemical power-stroke driven by ATP binding. They furthermore suggest that short-lived intermediates inaccessible to standard structural studies play a central role in active transport.

**Fig. 1.**
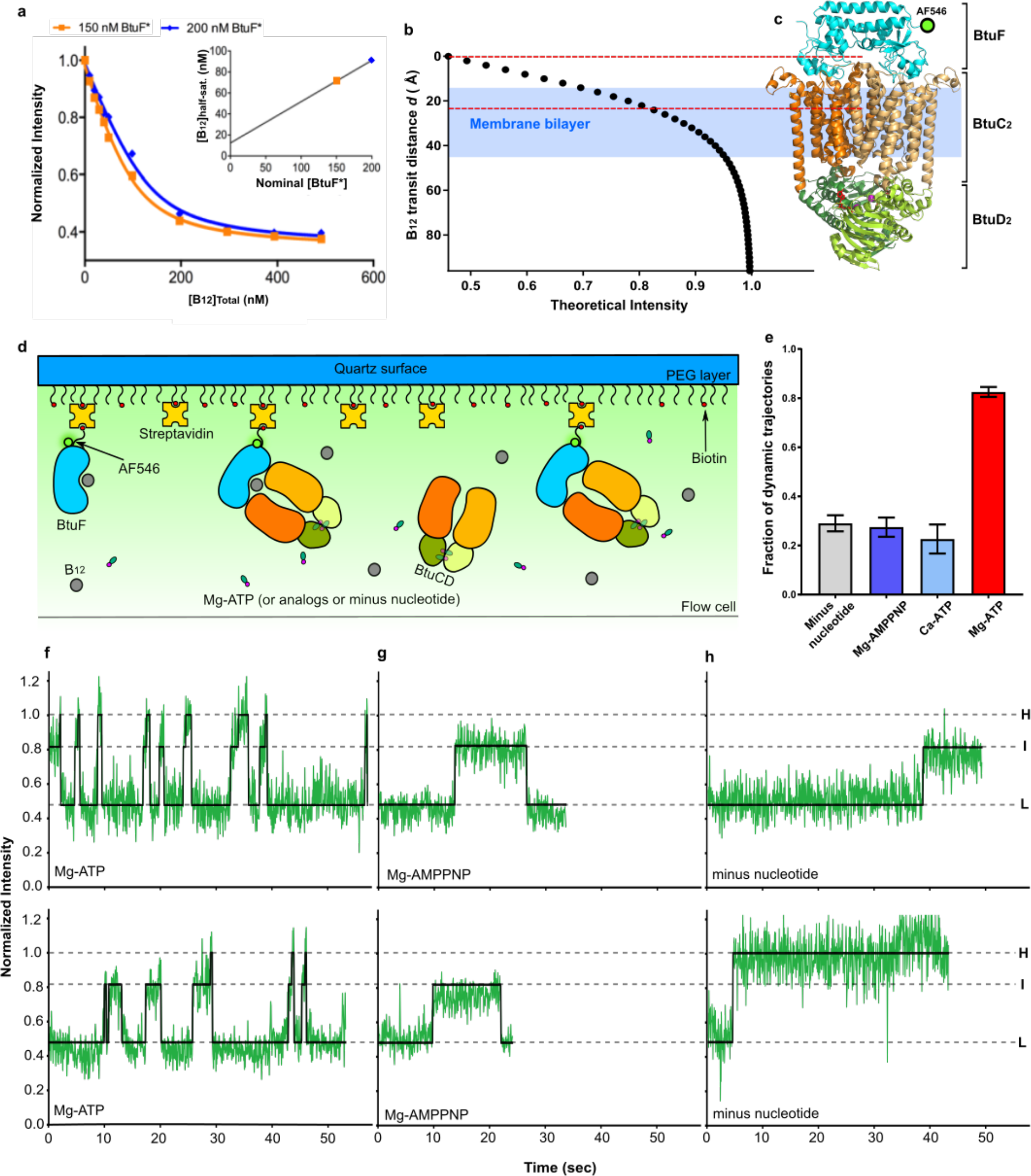
smFRET experimental setup and representative trajectories. **a**, Ensemble experiment titrating B_12_ onto BtuF* shown with global fits to the quadratic binding equation, which give *Kd* equal to 23 ± 6 nM. *Insert*, Plotting [B_12_] at half-saturation (determined by single-curve fitting) vs. nominal receptor concentration yields stoichiometry estimates of ∼0.8 and *K_d_* estimates of ∼12 nM. **b**, Theoretical fluorescence intensity level plotted against transit distance (*d*) of B_12_ from the binding site on BtuF* using the AF546 transition dipole moment (TDM) position from **Fig. S 1b** that matches the intermediate or I fluorescence state (E_FRET_ = 0.82) observed in the experiments reported below to the center of the internal cavity in BtuC in all available crystal structures. The systematic calibration analysis in **Fig. S1b** shows that the transit distance in the I state corresponds to approximately the middle of the membrane for all physically reasonable locations of the TDM. **c**, Structure of BtuCD-F complex (PDB 4FI3). The green circle represents labeling site of AF546. The blue background approximates transmembrane region. Red dash lines approximate binding sites of B_12_ in BtuF and BtuCD. **d**, Schematic of smFRET experiment setup. AF546-labeled BtuF (BtuF*) is immobilized on a passivated quartz surface by biotin-streptavidin interaction. During experiments, BtuF* is incubated with B_12_, nanodisc-BtuCD, and MgATP or nonhydrolyzable analogs. (See methods for details). **e**, Fractional number of dynamic trajectories found in experiments containing no nucleotides, 0.5 mM Mg-ATP, Ca-ATP, or Mg-ATP while maintaining same concentrations of BtuCD (1 µM) and B_12_ (1 µM). **f-h**, Representative dynamic trajectories in smFRET experiments summarized in panel **e**. Three intensity levels (L, I and H) are shown in dash lines. All smFRET experiments were conducted at 21 ± 1 °C.

**Fig. 2.**
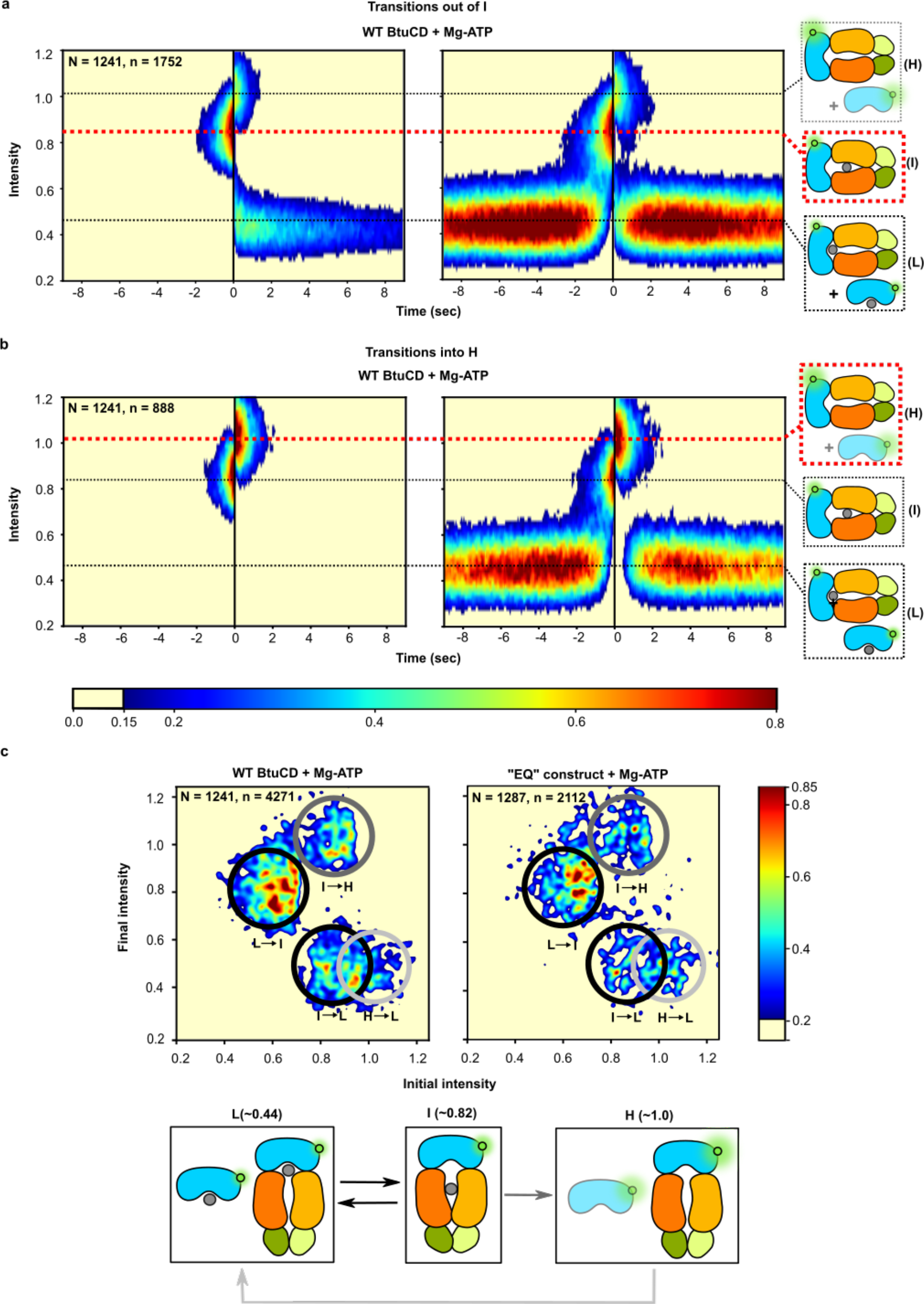
ATP hydrolysis drives unidirectional conformational transitions. **a**, 2D surface contour plots depicting the synchronized time evolution of smFRET-Q populations in an experiment conducted in the presence of 0.5 mM Mg-ATP, 1 µM BtuCD and 1 µM B_12_ (defined as the reference condition). Transitions out of the I (∼0.82) smFRET-Q state are aligned at the last datapoint preceding that transition (t = 0 in this plot). The relative populations of trajectories are color-coded as calibrated by the scale-bar below panel **b** (with values normalized to the 2D time/Intensity bin of maximum population). The plot on the left includes data from each trajectory only during the time window in which it is in the I state preceding the aligned transition or the new state immediately following that transition; this presentation highlights the final states produced by the transition and the lifetimes of the initial/final states preceding/following the transition. The plot on the right shows all data in a window of ± 9 sec around the transition point, which enables visualization of the complete reaction cycle. **b**, Synchronized plots for the same experiment with trajectories showing transitions into the H (∼1.0) smFRET-Q state aligned at the first datapoint following that transition (t = 0). The right panels in **a** and **b** show schematics of the complexes involved in the transitions; as discussed in the main text, *apo* BtuF* is not directly observed within the time resolution of these experiments (**Fig. S3b**). **c**, The left panel shows a transition density plot (TDP) for the aforementioned experiments conducted on the WT BtuCD construct in the reference condition. The relative populations are color-coded as calibrated by the scale-bar on the right with values normalized as described above. Black circles highlight transitions between L and I, while the dark gray circle highlights transitions from I to H, and the light gray circle highlights transitions from H to L. The right panel shows a TDP for an equivalent experiment conducted with the tandem “EQ” BtuCD construct illustrated in **Fig. S10**; it is shown here for direct comparison to the data from the WT construct. The bottom panel schematizes the transitions observed in the TDPs. In all panels, “N” gives the total number of trajectories in the experiment, while "n" gives the number of synchronized transitions (panels **a-b**) or the total number of transitions (panel **c**).

Several specific features of the BtuCD-F transport mechanism remain enigmatic. One is that, in the absence of both B_12_ and ATP, BtuF rapidly forms a hyper-stable (locked) complex with BtuCD that can only be destabilized by the presence of ATP^14,15^, which partially contributes to its high basal ATPase activity22. However, no comparable phenomenon involving apparently futile ATP hydrolysis has been observed for type I ABC importers^31–33^, which generally show tight coupling of hydrolysis to transport. In our ensemble fluorescence spectroscopy study, we showed that ATP hydrolysis, instead of ATP binding, leads to dissociation of the locked BtuCD-F complex^34^. We also estimated that the activation enthalpy of the locking reaction to be ∼103 kJ/mol, which suggests large conformational changes in the BtuCD-F complex during the reaction. These results lead to the hypothesis that the locking reaction plays an essential role in transport of B_12_^34^.

It is also unclear how ATP hydrolysis is coordinated between the two active sites in the BtuCD homodimer. Similar to many ABC proteins, the ATPase activity of BtuCD exhibits strong positive cooperativity (Hill coefficient ∼2)^16^, indicating its two active sites cooperate in binding ATP. While this property points towards a conformational reaction mechanism preserving the symmetry of the homodimer, there is no obvious physicochemical mechanism to trigger hydrolysis simultaneously at two spatially separated active sites, suggesting it occurs sequentially but with an unclear time interval between the hydrolysis reactions in the two active sites. A dynamic power stroke driven by hydrolysis of a single ATP would presumably push the BtuCD homodimer at least transiently into an asymmetric state^35^. This inference is consistent with the asymmetries observed in the structures of both BtuF21 and the locked BtuCD-F complex^17,18^. In our ensemble study, we also discovered functional asymmetry in the interaction between the two lobes of BtuF and the BtuC dimer34. Some ATPases that form higher-order oligomeric rings are known to employ sequential reaction mechanisms involving progression through structurally and functionally asymmetrical conformations^**36**^, but symmetrical homodimers are generally assumed to employ symmetrical conformational reaction mechanisms involving coordinated enzyme turnover at the interacting active sites^4,29,37–42^. However, mechanistic details of this kind remain obscure for most enzyme families given the challenges involved in characterizing dynamic conformational changes using standard structural methods.

Addressing such mechanistic questions effectively requires characterization of short-lived intermediates. We therefore designed a fluorescence reporter system to monitor movement of B_12_ through single BtuCD-F complexes on an ∼100-millisecond time scale. Using this system, we were able to visualize in real time, and establish the kinetics of, the complete ATP-powered transport cycle in nanodisc-reconstituted BtuCD-F, including binding of B_12_ to BtuF, initial movement of B_12_ into the TMD of BtuCD-F, release of B_12_ from the TMD, and return of the transporter to the conformational ground state. These data establish a novel transport mechanism that rationalizes the observations reported above. The homodimeric BtuCD assembly undergoes asymmetrical conformational changes in which three sequential mechanochemical power strokes drive progressive vectorial translocation of B_12_ across the membrane, with the first and last each being dynamically driven by hydrolysis of a single ATP molecule and the middle one being driven by the energy of BtuF binding.

## A molecular ruler to monitor vitamin B_12_ movement during transport

The visible emission spectrum of Alexa Fluor 546 (AF546) overlaps the absorption spectrum of B_12_, which produces distance-dependent FRET-induced quenching (FRET-Q) of AF5461923. We previously labeled BtuF with AF546 (BtuF*) to create a fluorescence reporter for its interaction with B_12_, which we used to characterize the mechanistic details of the BtuF*-BtuCD locking reaction summarized above^34^. We herein exploit that previously characterized system to report in real time on the location of B_12_ during transport. **Fig 1** summarizes the essential features of the FRET-Q system for the experiments reported in the current paper, and we employed a related approach to verify that our purified BtuCD mediates highly efficient, ATP-driven extrusion of B_12_ when reconstituted into sealed phospholipid vesicles (**Fig. S2**). At saturation, binding of B_12_ quenches BtuF* intensity by ∼54% in ensemble FRET-Q experiments in the absence of BtuCD (**Fig. 1a**), and fitting of fluorescence quenching curves^34^ (*e.g.*, as shown in **Fig. 1a**) yields a dissociation constant (*K_d_*) of ∼7.9 nM for the BtuF*•B_12_ binding reaction, matching previous reports^19,34,43^. During the transport process, B_12_ moves away from AF546-labeled BtuF*, which reduces AF546 quenching; the resulting increase in AF546 intensity, which we call “dequenching” provides a molecular ruler to monitor B_12_ movement through BtuCD during transport (**Fig. 1b-c** and **Fig. S1b**). We harnessed this FRET-Q-based photophysical ruler to develop a system to monitor the translocation of B_12_ through single transporters in real time, and these “smFRET-Q” measurements have enabled us to observe and characterize the transient processes that occur during ATP-driven transport of B_12_ through the BtuCD-F* complex that are obscured in ensemble experiments, which average signals across the protein population.

## Observation of ATP-driven B_12_ transport dynamics using smFRET-Q

To image individual BtuF* molecules using total internal reflection fluorescence (TIRF) microscopy, BtuF* was biotinylated and tethered to the polyethylene glycol-silane (PEG-silane)/biotin-PEG-silane-derivatized, streptavidin-treated surface of a quartz microfluidic flow-cell through a biotin-streptavidin-biotin bridge (**Fig. 1d**). Movement of B_12_ away from its binding site on individual surface-tethered BtuF* molecules is monitored via distance-dependent dequenching of the fluorescence of the AF546 covalently bonded to BtuF* (**Fig. 1b-c**). The intensity *versus* time trajectories recorded for individual BtuF* molecules (**Fig. 1f-h**) were normalized to the initial intensity in each trajectory prior to addition of B_12_, and the relative intensity calculated in this manner was used to quantify smFRET-Q levels.

Upon introducing a sub-saturating concentration (10 nM) of B_12_ into the flow-cell in the absence of BtuCD, the trajectories began to fluctuate between a ∼0.46 relative intensity state corresponding to ∼54% quenching and a ∼1.0 relative intensity state corresponding to ∼0% quenching (**Fig. S3a**). Based on the close match of these values to those in our ensemble FRET experiments^34^ (**Fig. 1a**), we interpreted the ∼0.46 state to be the B_12_-bound state of BtuF*(BtuF*•B_12_), the ∼1.0 state to be the B_12_-free state (apo BtuF*), and the transitions between them to reflect spontaneous binding and release of B_12_ by BtuF*. Kinetic analysis of the trajectories at 10 nM B_12_ using a 2-state Hidden Markov Model (HMM) yields rate constants of *k_binding_* equal to 0.065 ± 0.005 sec-1•nM-1 and *k_release_* equal to 0.130 ± 0.010 sec-1. The corresponding 2 nM *K_d_* (given by the ratio of *k_release_* /*k_binding_*) represents a reasonable match^44,45 44,45^ to the value of ∼8 µM given by ensemble experiments conducting using several methods^19,34,43^ (**Fig. 1a**).

The measured rates indicate that, at 1.0 µM concentration, B_12_ binds to BtuF* at a rate of 65 sec-1, significantly faster than the 16.6 sec-1 data acquisition rate of our detector (reciprocal of the 60 msec integration time). Therefore, at this B_12_ concentration, release of B_12_ from BtuF* should be followed by rebinding within the time interval used to collect a single data point, implying the emission intensity of individual BtuF* molecules should remain constant in the 0.46 state in trajectories acquired in the presence of 1.0 µM B_12_. This inference is confirmed in **Fig. S3b**. Therefore, smFRET-Q experiments to characterize transport through BtuCD were performed at the saturating, 1.0 µM concentration of B_12_ so that fluctuations in the relative intensity trajectories would overwhelmingly represent movement of B_12_ through BtuCD-F* rather than cycles of B_12_ binding and release by BtuF*.

Simultaneous introduction of 1.0 µM nanodisc-reconstituted BtuCD and 0.5 mM Mg•ATP in addition to 1.0 µM B_12_ in smFRET-Q experiments resulted in trajectories that exhibited ATP-hydrolysis-dependent fluctuations between three distinguishable relative intensity states (**Fig. 1e-h**). We hypothesized that these that these fluctuations, which occur at a much higher rate in the presence of Mg•ATP than in the absence of nucleotide or the presence of non-hydrolyzable analogs (Fig. 1e), reflect progressive movement of B_12_ away from BtuF* during the transport cycle. Based on their mean values, we designated the three observed relative intensity states as Low (L, 0.44 ± 0.01), Intermediate (I, 0.82 ± 0.03), and High (H, 1.02 ± 0.03) (**Figs. 1f-h** **& 2**). The ∼56% quenching in the L state matches the level observed in our ensemble FRET-Q34 and smFRET-Q experiments upon binding of B_12_ to BtuF* in the absence of BtuCD (**Fig 1a** and **Fig. S3**). Based on our ensemble experiments simultaneously measuring FRET-Q and anisotropy^34^, the L state represents a dynamic equilibrium between free BtuF*•B_12_ and the pre-transport BtuF*•B_12_•BtuCD complex prior to movement of B_12_ into BtuCD (states 1-2 on the left in **Fig. 4** below).

The ∼18% quenching in the I state is consistent with an ∼25 Å movement of B_12_ away from its binding site in BtuF* during the Mg•ATP-driven reaction cycle (**Fig. 1b-c**). Notably, this distance coincides with the location of a previously reported cavity within the TMDs in BtuCD12,13,24 (**Fig. S1a**) based on systematic analysis of all physically plausible AF546 locations (**Fig. S1b**).

The ∼0% quenching in the H state indicates that the B_12_ transport substrate has moved outside the range of detectable FRET from the AF546 fluorophore covalently bound to BtuF* (**Fig. 1b**). This state must have BtuF* sequestered in a complex that makes its B_12_ binding site inaccessible because, as explained above, the lifetime of *apo* BtuF* prior to rebinding B_12_ is too short to be captured by our detector in the presence of the 1.0 µM B_12_ concentration used in these experiments (**Fig. S3b**). Our ensemble FRET-Q experiments^34^ demonstrate that the B_12_ binding site in BtuF* is inaccessible in the locked BtuCD-F* complex that can only be dissociated by Mg•ATP hydrolysis. Therefore, we infer that formation of the H state involves movement of B_12_ away from a cavity in the TMD in BtuCD concomitant with sealing of BtuF* on the periplasmic surface of BtuCD to form the locked BtuCD-F* complex. Given that BtuF* locking sterically blocks B_12_ exit from the periplasmic side of the transporter, formation of the H state most likely involves exit of B_12_ from the distal cytoplasmic side of the transporter to complete the B_12_ translocation process. The efficient TM transport of B_12_ mediated by our purified proteins reconstituted into liposomes (**Fig. S2**) and detailed kinetic analyses of smFRET-Q data in nanodiscs (**Figs. 2**-**3** below) support this interpretation.

**Fig. 3.**
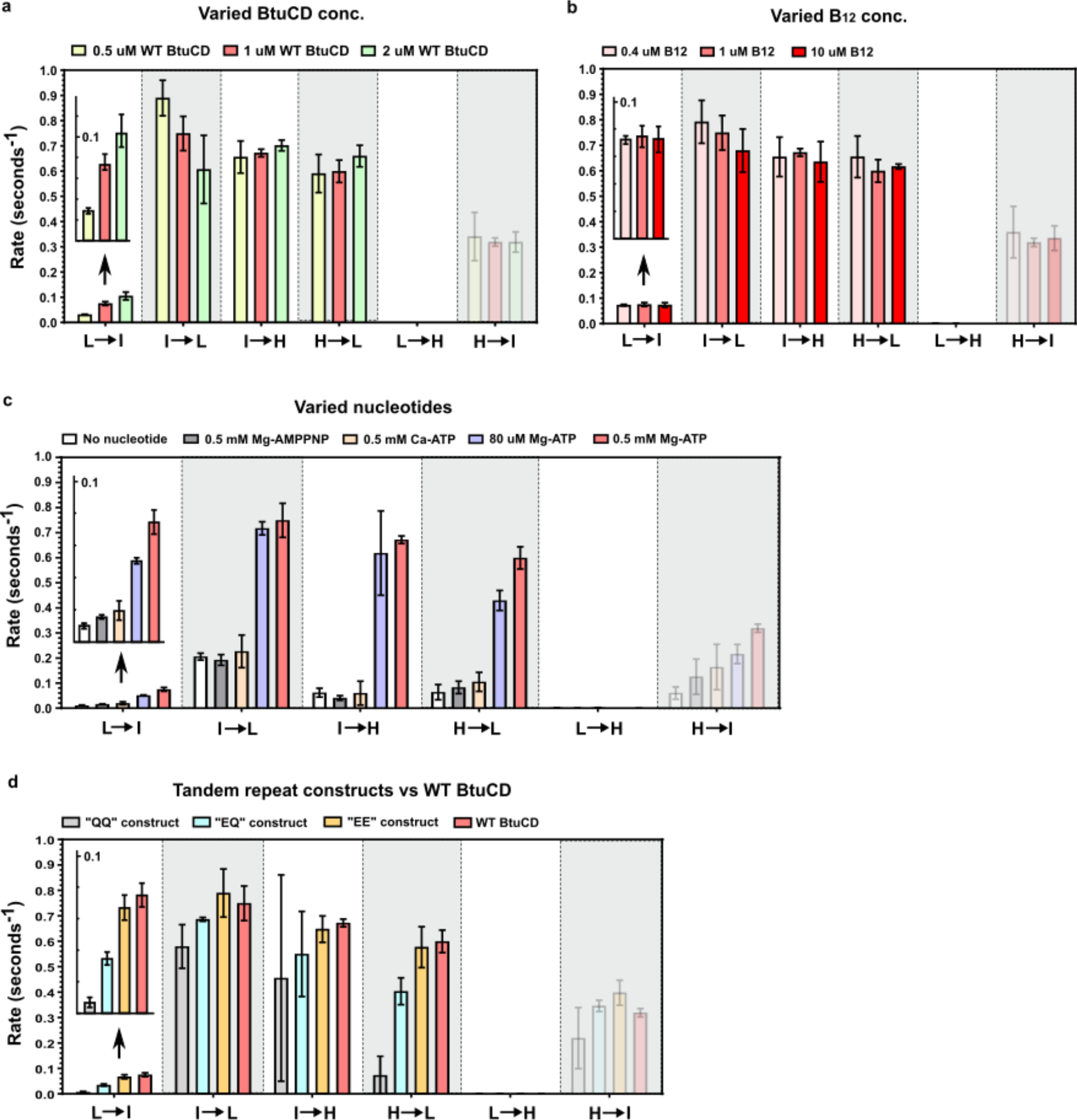
Kinetics of conformational transitions. **a**, Effects of BtuCD concentrations on kinetics. Experiments maintain the same concentrations of B_12_ (1 µM) and Mg-ATP (0.5 mM) but vary the concentrations of BtuCD (0.5 µM, 1 µM and 2 µM). **b**, Effects of varied B_12_ concentrations on kinetics. Experiments maintain the same concentrations of BtuCD (1 µM) and Mg-ATP (0.5 mM) but vary the concentrations of B_12_ (0.4 µM, 1 µM and 10 µM). **c**, Effects of ATP binding and hydrolysis on kinetics. Experiments maintain the same concentrations of BtuCD (1 µM) and B_12_ (1 µM) but vary the nucleotide conditions: minus nucleotide, 0.5mM Mg-AMPPNP, 0.5mM Ca-ATP, 80uM Mg-ATP, and 0.5mM Mg-ATP. **d**, Effect of single ATP hydrolysis on kinetics. Experiments maintain the same concentrations of B_12_ (1 µM) and Mg-ATP (0.5 mM) but use different constructs of BtuCD at 1 µM: “QQ”, “EQ”, “EE” and WT. Error bars represent standard deviations obtained from triplicate experiments. L à H and Hà I transitions are shown in semitransparent bars due to the very low number of observed transitions.

**Fig. 4.**
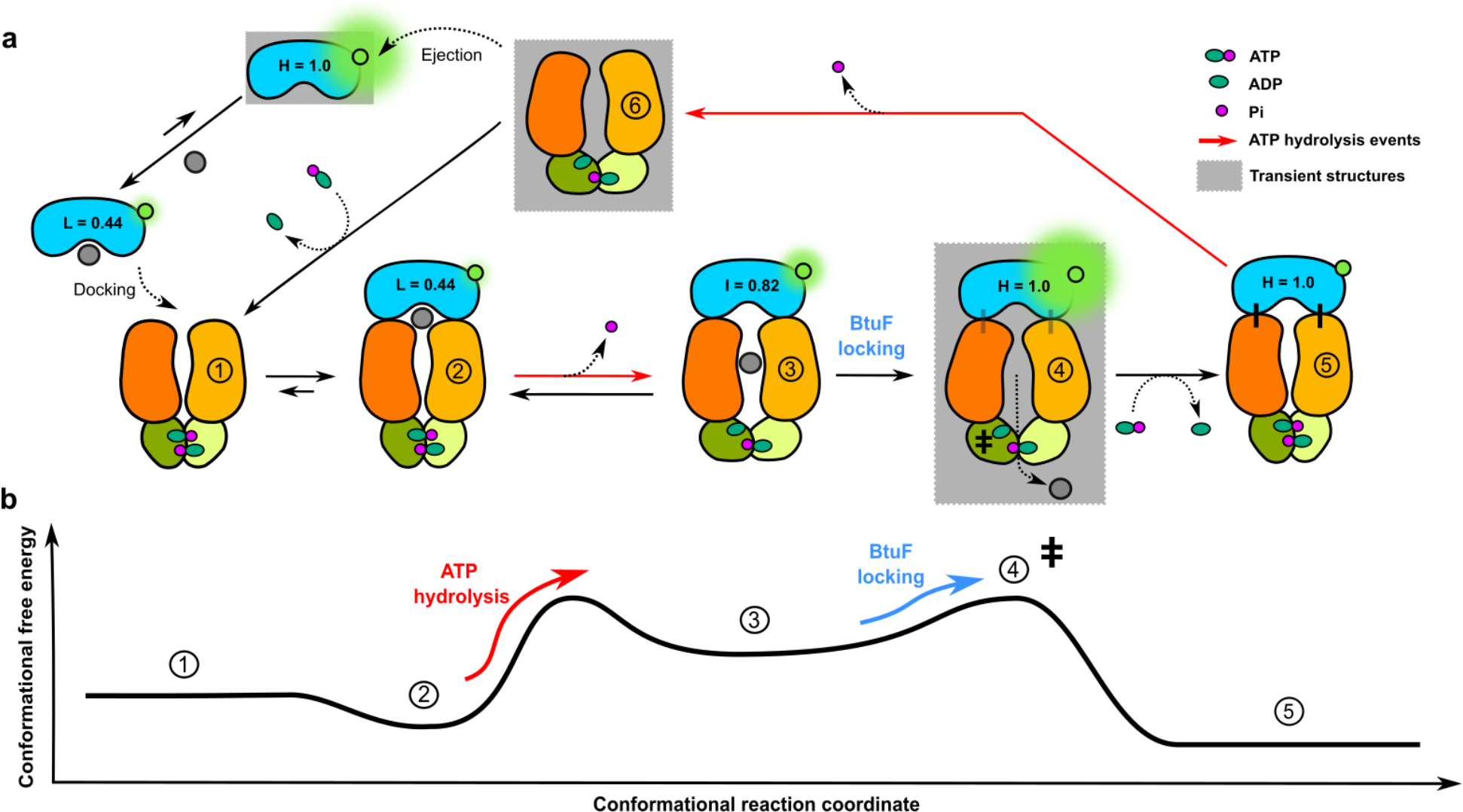
Proposed mechanism and energy diagram. **a**, Schematic showing the inferred BtuCD-F* transport mechanism. BtuF* exhibits three smFRET-Q states (L, I and H) while BtuCD cycles through six conformational states (1 to 6). State 1 represents Mg-ATP-bound BtuCD ready to bind to B_12_-loaded BtuF*. State 2 represents initial docking of B_12_-loaded BtuF* to BtuCD. State 3 represents BtuCD with B_12_ residing within its TMDs after hydrolysis of the first ATP molecule. State 4 represents a transient inward-facing BtuCD conformation leading to the exit of B_12_ at the cytoplasmic side of the transporter. State 5 represents locked hyper-stable BtuCD-F* complex that prevents rebinding of B_12_. Hydrolysis of a second ATP molecule results in ejection of BtuF* from BtuCD in state 6. Transient structures with lifetimes shorter than the60 msec frame time of our detector are shown in gray background. **b**, Corresponding conformational free energies.

To begin dissecting the role of Mg•ATP binding *vs.* hydrolysis in the transport cycle, we recorded intensity trajectories maintaining the same constant and saturating 1.0 µM concentrations of B_12_ and BtuCD but either omitting Mg•ATP (minus nucleotide — **Fig. 1h**) or substituting it with one of two non-hydrolysable analogs (Mg•AMPPNP or Ca•ATP — **Fig. 1g** and data not shown, respectively). These experiments show a very large decrease in transition rates between all intensity states in the presence of the non-hydrolyzable ATP analogs (*e.g.*, as shown in **Fig. 1g** *vs.* **1f** and systematically quantified in **Fig. 3c** below). The dramatically more frequent transitions between states observed in the presence of Mg•ATP (**Fig. 1e**) suggest they are driven by cycles of Mg•ATP binding and hydrolysis by the BtuCD-F* transporter complex. Consistent with this inference, 2-color smFRET experiments using a single-site on each BtuC subunit stochastically labeled with a Cy3 donor and Cy5 acceptor show that there is no significant change in the conformations adopted by either BtuCD or BtuCD-F upon addition of Mg•AMPPNP (**Figs. S4-S6**). In contrast, a population with a significantly wider separation between the TM *α*-helices at the cytoplasmic surface of BtuCD appears exclusively in the presence of Mg•ATP but not in the absence of nucleotide or in the presence of Mg•AMPPNP. This state represents ∼4% of observed E_FRET_ values in trajectories measured in the presence of Mg•ATP without BtuF and ∼6% of those observed in the presence of both Mg•ATP and BtuF. These observations suggest that ATP hydrolysis drives opening of the cytoplasmic gate of the internal cavity in BtuCD, which is ultimately needed for B_12_ transport.

## Unidirectional progression through conformational states driven by ATP hydrolysis

We evaluated the time evolution of progression through the L, I, and H relative intensity states using two analyses that harness a global HMM optimized to describe the smFRET-Q trajectories. The first employed “synchronized” 2D surface contour plots, which superimpose all individual trajectories collected under a single condition after aligning them at the transition time into or out of one specific intensity state (**Fig. 2a-b** **and Fig. S6**). The second employed transition density plots (TDPs) (**Fig. 2c** and **Fig. S7**), which systematically quantify the frequency of transitions between the relative intensity states. Both analyses show that the progression through the I → H → L states is unidirectional rather than reversible.

Therefore, this transition sequence represents a non-equilibrium process, consistent with expectations for an irreversible ATP-driven active transport process.

**Fig. 2a** shows surface-contour plots synchronized at the time of transition out of the I relative intensity state to visualize the time evolution of the population of BtuCD-F*•B_12_ transporter complexes as they leave this state. These plots demonstrate that, in the presence of Mg•ATP, BtuCD-F* progresses through two alternative transition cycles with high frequency: L → I → L and L → I → H → L, both of which show a much longer dwell-time in the L relative intensity state than the I or H states. **Fig. 2b** shows surface-contour plots synchronized instead at the time of transition into the H relative intensity state to visualize the time evolution of molecules as they enter that state. These plots demonstrate that the transitions through the H state occur exclusively via the unidirectional sequence I → H → L. The unidirectionality of this transition sequence is confirmed by the striking asymmetry across the diagonal of the TDP for the experiment conducted in the presence of Mg•ATP, which shows a high frequency for the I → H transition compared to a negligible frequency of the reverse transition (**Fig. 2c**). Importantly, the first two transitions in the L → I → H → L cycle, which is observed in the presence of MgATP, show a monotonic decrease in BtuF* quenching, indicating progressive B_12_ movement away from the fluorophore covalently linked to BtuF*, consistent with transport of B_12_ through BtuCD-F*.

During the irreversible I → H transition, B_12_ moves from a location inside the TMDs of the transporter to a location outside the range of FRET to the AF546 fluorophore on BtuF* in less than the 60 msec time window for a data point. Given that the kinetically competing I → L transition represents return of B_12_ from a location in the TMD of the transporter to its original location bound to BtuF* on the periplasmic side, the I → H transition is most parsimoniously explained by release of B_12_ on the cytoplasmic side of the membrane coupled to locking of BtuF* to BtuCD on the periplasmic side of the membrane, which blocks the periplasmic exit channel from the TMDs. As explained above, formation of the locked BtuCD-F* complex must occur concomitantly with entry into the H relative intensity state in order to explain its sustained ∼1 sec occupancy (**Fig. 2b**) given that rebinding of B_12_ to *apo* BtuF* is substantially faster than the time of a single data point (**Fig. S3b**) in the presence of the saturating 1.0 µM concentration of B_12_ used in these experiments. Based on our ensemble study showing that the dissociation of the locked BtuCD-F* complex requires Mg•ATP hydrolysis^34^, we infer that the H → L transition involves ejection of BtuF* from the complex induced by Mg•ATP hydrolysis followed by rapid binding of B_12_ to the released *apo* BtuF*. Therefore, the L → I → H → L transition sequence is consistent with stepwise translocation of B_12_ from its initial, pre-transport binding site on BtuF* (the L state), to the cavity inside the TMDs of BtuCD (the I state), followed by release of B_12_ from the opposite side of the transporter (the H state), and finally binding of a new B_12_ molecule to BtuF* to return the system to the pre-transport configuration (the L state). (See mechanistic schematic in **Fig. 4a** below.)

Synchronized surface-contour plots (**Figs. S7-S8**) and TDPs (**Fig. S9a-c**) constructed using the intensity trajectories recorded in the absence of ATP or the presence of non-hydrolyzable analogs (**Fig. 1g-h**) show dramatically different transition patterns in addition to greatly reduced transition frequencies (**Fig. 1e**). The I state has a systematically lower intensity than the I state observed in the presence of Mg•ATP (∼0.75 *vs.* ∼0.82; **Fig. 2a-b** *vs.* **Fig. S7**, summarized in **Fig. S10**), leading us to designate it as the I’ relative intensity state. The experiments in the absence of ATP or the presence of non-hydrolyzable analogs show almost exclusively a reversible L → I’ → L transition sequence (reflected in diagonal symmetry in the TDPs in **Figs. S9a-c**). The I’ state observed in these experiments, and the L → I’ → L transition sequence shows substantially longer dwell times in both the L and I’ states compared the L and I states observed in the presence of Mg•ATP (shown qualitatively in **Fig. 1f-h** and **Fig. S7a** and quantified in **Fig. 3b** below). The lower intensity of the I’ state indicates that B_12_ does not move as far into the TMDs of BtuCD-F* during its slower excursions when there is no nucleotide bound or when non-hydrolyzable Mg•AMPNP or Ca•ATP are bound compared to its much faster movement into the TM cavity when Mg•ATP is bound and being hydrolyzed. These observations strongly support the conclusion that Mg•ATP hydrolysis drives translocation of B_12_ from its initial binding position on BtuF* in the L state into the TMDs of the BtuCD-F* complex in the I state.

## Kinetic analyses of relative intensity trajectories to elucidate mechanistic details of the transport cycle

We used HMM-based kinetic analyses to quantify the rates of transitions between the L, I, and H states observed in additional experiments in which we varied the concentrations of BtuCD, Mg•ATP, or B_12_ (**Table S1** in the ***Supplementary Information***). These experiments demonstrate that the rate of the L → I transition is reduced when BtuCD concentration is reduced, while the rates of all the other transitions remain unchanged (**Fig. 3a**). These kinetic observations are consistent with BtuF*•B_12_ binding to BtuCD prior to or during the L → I transition in which B_12_ moves from its binding site on BtuF* into the TMD of BtuCD (**Fig. 4**). They moreover suggest that the resulting BtuCD-F* complex in the I state must remain associated continuously without exchange of any of the protein components prior to completion of the subsequent I → L transition or I → H → L transition sequence because the rates of these transitions are unaffected by variation in BtuCD concentration.

No changes are observed in the rates of any of the transitions when B_12_ concentration is varied over a 25-fold range in which its rate of rebinding to *apo* BtuF* remains higher than the data acquisition rate of our detector (**Fig. 3b**). The observation that neither BtuCD nor B_12_ concentration influences the rates of the two kinetically competing transitions out of the I state (I → L and I → H) reinforces the conclusion presented above that B_12_ is sequestered inside the cavity in the TMDs of the BtuCD-F* complex following the L → I transition in which Mg•ATP hydrolysis induces B_12_ movement into this cavity from its initial binding site on BtuF* (*i.e.*, as schematized in the transition between states 2 → 3 in **Fig. 4a**). The reverse I → L transition can represent either B_12_ movement from that cavity directly back to its original binding site on BtuF* or release of BtuCD followed by rapid rebinding of a B_12_ molecule from bulk solution back to the surface-tethered BtuF*. The I → H transition must represent locking of BtuF* onto the periplasmic surface of BtuCD simultaneously with the movement of B_12_ out of the cavity inside its TMDs because, except in the locked state^34^, BtuF* will rapidly rebind B_12_ and return to the L state in less time than the duration of a single data point under the saturating 1 µM B_12_ concentration used in these experiments (**Fig. S3b**). Because locking of BtuF* on the periplasmic surface blocks the exit pathway to the periplasm from the central cavity in BtuCD, release of B_12_ simultaneously with locking strongly supports the conclusion that the I → H transition represents release of B_12_ on the cytoplasmic side of the complex to complete transport. The simultaneity of these processes furthermore supports our earlier inference that locking plays a direct role in the transport process^34^.

The kinetic analyses furthermore demonstrate that the rates of both the L → I and H → L transitions are reduced when Mg•ATP concentration is reduced, while the rates of the other transitions do not change (**Fig. 3c**). Combined with the observation that the L → I transition occurs rapidly in the presence of Mg•ATP but comparatively very slowly in the absence of nucleotide or the presence of non-hydrolyzable analogs (**Figs. 1f-h and 3c)**, the reduction in the rate of this transition at lower Mg•ATP concentration is consistent with a requirement for Mg•ATP binding and hydrolysis by BtuCD for the movement of B_12_ from its binding site on BtuF* in the L state into the TMD of BtuCD in the I state (*i.e.*, the transition between states 2 → 3 in **Fig. 4a**). Furthermore, the reduction in the rate of the H → L transition at lower Mg•ATP concentration implies that a second Mg•ATP binding and hydrolysis event is required for this transition (between states 5 → 6 in **Fig. 4a**), and this observation strongly supports our inference that it represents Mg•ATP hydrolysis driving dissociation of the locked BtuCD-F* complex formed after BtuF* drives a conformational transition that releases B_12_ from the TMD in BtuC to the cytoplasm during the intervening I → H transition (between states 3 → 5 in **Fig. 4**).

### Ensemble and smFRET-Q experiments on covalently linked, tandem BtuD_2_ constructs

To characterize the stoichiometry of Mg•ATP hydrolysis and coordination between the two ATPase active sites in the homodimeric BtuCD complex, we employed a variant of a previously reported construct containing two covalently linked BtuD molecules16. This tandem BtuD_2_ construct enables glutamate-to-glutamine (E159Q) mutations that preserve ATP-binding affinity but block ATP hydrolysis17 to be introduced into the catalytic base in either one or both of the ATPase active sites in the transporter complex (**Fig. 3d** and **Fig. S10a**). Compared to the previously reported BtuD_2_ construct, which exhibited slightly impaired function16, we lengthened the inter-BtuD linker from (G4S)4 to (G4S)^5^. In addition to a control construct with two fully wild-type (WT) ATPase active sites, we made equivalent constructs introducing the hydrolysis-blocking E159Q mutation into one or both of the ATPase active sites (**Fig. S10a**). We call these BtuC-BtuC-BtuD_2_ complexes ‘EE’, ‘EQ’, or ‘QQ’ based on whether there are zero, one, or two E159Q mutations, respectively, in the tandem BtuD_2_ domains.

Our EE construct behaves quantitatively equivalently to WT BtuCD in Michaelis-Menten kinetic assays of ATPase activity (**Fig. S10b**), ensemble FRET and anisotropy assays (**Fig. S10c**), and single-molecule transport assays (**Figs. 2c** and **3d** and **Figs. S7 & S9d-f**), except for a reduction in the cooperativity of the ATPase activity that is observed in the Michaelis-Menten kinetic assays (**Fig. S10b**). The very similar values for the Michaelis constant (*K_M_*) and maximum velocity (*V*_max_) obtained from the kinetic analyses of the EE construct *vs.* WT BtuCD suggest that the reduced cooperativity of EE reflects a pre-configuration of the ATPase active site into a high-affinity conformation for Mg•ATP binding due to a spatial constraint introduced by the linker.

The EQ and QQ constructs also behave equivalently to WT BtuCD in ensemble FRET and anisotropy experiments prior to Mg•ATP addition (**Fig. S10c**). However, they reduce the *V*max for steady-state ATPase activity to 38% and an undetectable level (**Fig. S10b**), respectively, confirming the glutamate-to-glutamine mutations block ATP hydrolysis. The EQ transporter complex also behaves qualitatively equivalently to the WT BtuCD in smFRET-Q experiments (**Fig. 2c** and **Figs. S7 & 9e**) but results in relative intensity trajectories that exhibit altered rate constants for some transitions (**Fig. 3d**), as discussed below. In contrast, trajectories recorded from the QQ construct show essentially no transitions (**Fig. 3d** and **Figs. S7** and S**9f**) even through it interacts with BtuF* equivalently to WT BtuCD prior to Mg•ATP addition in ensemble experiments (**Fig. S10c**).

These observations reinforce the conclusion that ATP hydrolysis is required for transport of B_12_ from its binding site on BtuF* into the TMD cavity in BtuCD during the L → I transition. Kinetic analyses of the relative intensity trajectories recorded for the EQ complex (**Fig. 3d**) strongly support sequential hydrolysis of one Mg•ATP molecule during the L → I transition and a second during the subsequent H → L transition that resets the transporter to the initial state (states 2 → 5 in **Fig. 4**). The rates of these two transitions at a saturating Mg•ATP concentration are both significantly reduced in the EQ construct compared to the EE construct and WT BtuCD, while the rates of the other transitions remain unchanged (**Fig. 3d**). In the EQ construct with only one of the two Mg•ATP binding sites capable of performing hydrolysis, the rate of the L → I transition is reduced by 50% compared to the EE or WT constructs (**Fig. 3d**). However, the rates of the ensuing I → L and I → H transitions are unchanged in the EQ construct compared to WT (**Fig. 3d**). These observations imply that only one of the two bound Mg•ATP molecules is hydrolyzed during the L → I transition that moves B_12_ from its binding site on BtuF* into the TMD cavity in BtuCD and that the second Mg•ATP molecule remains bound without being hydrolyzed during either of the ensuing I → L (states 3 → 2 in **Fig. 4**) or I → H transitions (states 3 → 5 in **Fig. 4**).

The observation that the rate of the H → L transition depends on Mg•ATP concentration while the rate of the preceding I → H transition does not (**Fig. 3c**) indicates that the Mg•ATP binding step in the Mg•ADP/Mg•ATP exchange process occurs after formation of the locked BtuCD-F* complex in the H state (*i.e.*, between states 4 → 5 in **Fig. 4**), but before BtuF* dissociation during the H → L transition (states 5 → 6 in **Fig. 4**). This transition occurs in a qualitatively similar manner in the EQ construct but at 70% of the rate of the EE or WT constructs, which strongly supports hydrolysis of a single Mg•ATP molecule driving BtuF* dissociation because the transition would be completely inhibited in a construct with one inactivated ATPase site if it required hydrolysis of both Mg•ATP molecules. The 30% reduction in the rate of the H → L transition in the EQ construct compared to WT is statistically significant given the uncertainties in our measurements (*p* = 0.008), but it is not significantly different from half the WT rate (*p* = 0.035), making it unclear whether the presence of a single E159Q mutation in the tandem BtuD_2_ construct modestly accelerates hydrolysis at the unmutated active site. Our experiments with tandem BtuD_2_ constructs therefore demonstrate that TM translocation requires two sequential Mg•ATP hydrolysis reactions, the first driving B_12_ movement from BtuF* into a cavity in the TMD of BtuCD (states 2 → 3 in **Fig. 4d**) and the second driving dissociation (states 5 → 6 in **Fig. 4d**) of the hyper-stable BtuCD-F* complex formed during release of B_12_ from that cavity to the opposite side of the transporter (states 3 → 5 in **Fig. 4**).

## Conclusions

The results presented here demonstrate that FRET from a donor-labeled transporter to a transport substrate that can act as an acceptor can be used in smFRET experiments to monitor the physical translocation of a substrate through a TM transporter in real time. We employ mFRET quenching (smFRET-Q) experiments (**Fig. 1-3** and **Figs. S3** & **S6**-**S7**) supported by ensemble Michalis-Menten (**Fig. S10b**), ensemble FRET-Q, and anisotropy experiments^34^ to establish a novel molecular mechanism (**Fig. 4**) in which two sequential Mg•ATP hydrolysis reactions, rather than Mg•ATP binding, dynamically drive stepwise protein conformational changes that mediate unidirectional translocation of vitamin B_12_ through *E. coli* BtuCD-F, a model type II ABC importer. The short lifetimes (< 1 sec) of the high-energy conformational states demonstrated by our smFRET-Q experiments to mediate transport (**Fig. 2a-b** and) make them inaccessible to conventional structural studies, explaining why extensive structural studies of BtuCD-F have not succeeded in elucidating its transport mechanism.

Following binding of B_12_ to *apo* BtuF, the resulting BtuF•B_12_ complex reversibly binds to the periplasmic side of BtuCD (states 1 → 2 in **Fig. 4**, corresponding to the L relative intensity level). In the absence of ATP or presence of non-hydrolyzable analogs, the B_12_ molecule in this docking complex slowly moves back-and-forth to a position inside the TMD of BtuCD approximately in the outer leaflet of the cytoplasmic membrane (corresponding to the I’ relative intensity level), but B_12_ exclusively moves back to its initial binding site on BtuF from this position (**External Data Figs. 7-9**). Hydrolysis of one Mg•ATP molecule by the initial docking complex drives a first mechanochemical power-stroke producing an ∼25 Å movement of B_12_ from its initial binding site on BtuF* to a position further inside the TMD in BtuCD (states 2 → 3 in **Fig. 4**, corresponding to the I relative intensity level), roughly matching the site of an internal cavity in the transporter in the available crystal structures (**Fig. 1b** and **External Data** **Fig. 1a**). The resulting kinetically sequestered BtuCD-F complex effectively represents a high-energy mechanical transition state for the B_12_ translocation reaction given its short lifetime (∼1.7 sec-1, **Figs. 2a-b** and **3**) combined with the observation that it partitions in a roughly 1:1 ratio between reversion to the initial BtuCD/BtuF•B_12_ docking complex (states 3 → 2 in **Fig. 4**) and release of B_12_ on the opposite cytoplasmic side of the transporter to complete the translocation reaction (states 3 → 5 in **Fig. 4**). The latter productive translocation reaction produces a hyper-stable locked BtuCD-F complex (corresponding to the H relative intensity level) that requires Mg•ADP/Mg•ATP exchange followed by hydrolysis of one additional Mg•ATP molecule to drive a second mechanochemical power-stroke that induces BtuF release to return the transporter to its starting conformation (states 5 → 6 in **Fig. 4** in **Fig. 4**).

The energy of formation of the locked complex, which we previously estimated at ∼103 kJ/mol^34^, likely helps drive the conformational change inducing B_12_ release to the cytosolic side of the transporter (state 4 in **Fig. 4**). Using the energy of BtuF binding on the periplasmic side of the BtuCD to drive this critical conformational change will prevent backflow of the large B_12_ molecule to periplasmic side of the transporter, which could explain evolution of a mechanism in which, rather than directly driving transport, Mg•ATP hydrolysis drives dissociation of a protein complex that is formed to drive transport. This transport mechanism in which two temporally separated hydrolysis events drive sequential mechanochemical power-strokes in a cooperative ATPase solves a theoretical conundrum concerning the lack of a clear mechanism to coordinate hydrolysis at two spatially-separated ATPase active sites to enable Mg•ATP hydrolysis to drive a single mechanochemical power-stroke.

Michaelis-Menten kinetic analysis shows that Mg•ATP binds to WT BtuCD-F with ∼2-fold positive cooperativity (**Fig. S10b**), indicating hydrolysis takes place after binding of two Mg•ATP molecules, which are likely to be tightly encapsulated in interfacial active sites in a transporter complex with approximate 2-fold symmetry in the BtuCD subunits24. During hydrolysis of a single Mg•ATP molecule, cleavage of the phosphodiester bond between its *β*- and *γ*-phosphates will lead to local electrostatic repulsion that dynamically forces the BtuD subunits apart at that interfacial active site46 while Mg•ATP likely remains tightly encapsulated in the other one. Therefore, given that Mg•ADP/Mg•ATP exchange precedes the second of the two sequential hydrolysis reactions in the transport mechanism established by our smFRET-Q experiments, the first hydrolysis event (states 2 → 3 in **Fig. 4**) is likely to drive the BtuCD subunits into an asymmetrical conformation. A fundamentally asymmetrical conformational reaction cycle has previously been proposed for some ABC superfamily proteins but not yet verified in structural studies35,47–49. Additional research will be required to elucidate the structure of the high-energy BtuCD-F conformations induced by Mg•ATP hydrolysis that drive B_12_ translocation and BtuF dissociation.

## Acknowledgements

This work was supported by a RISE grant to JFH & RLG and graduate research fellowships to LZ, JK, and JER from Columbia University. LZ and CDK-T were supported in part by the US National Institutes of Health Training Program in Molecular Biophysics (T32-GM008281), and CDK-T was also supported in part by the Graduate Fellowship Program of the US Department of Energy Office of Science (DEAC05-06OR23100). Additional support was provided by grants from the US NIH-NIGMS to JFH (GM127883) and RLG (GM137608 and GM084288) and grants from the US Cystic Fibrosis Foundation to JFH (HUNT18G0 and HUNT20G0). We thank D.C. Rees for provision of the WT BtuCD expression plasmid, the members of the Hunt and Gonzalez labs for scientific advice, and E.C. Greene for a critical review of the manuscript.

## Author contributions

LZ, JK, KL, JER, NKK, RLG, and JFH conceived the project and designed the experiments, which were performed by LZ, JK, KL, JER, and analyzed by LZ, JK, KL, JER, CDK-T, RLG, and JFH. LZ, JFH, and RLG drafted the manuscript and finalized it with help from all authors.

## Conflict of interest

JFH is a member of the Scientific Advisory Board of Nexomics Biosciences, Inc., and a consultant for Cyrus Biotechnology. The other authors declare no competing financial interests.

Note: The **Supplementary Information** for this manuscript contains 11 figures and 1 table.

## METHODS

### Plasmids and mutagenesis

A cysteine residue was appended onto the C-terminus of BtuF for fluorophore labeling via sulfhydryl-maleimide chemistry for both ensemble and smFRET experiments that measure fluorescent signal. It was followed by a hexahistidine (His_6_)-tag. Non-interacting BtuF (niBtuF – **Supplementary Information Figure 2**) was created by mutating residues Glu-72 and Glu-202 to alanine to block BtuCD binding34 in order to create a biosensor for B_12_ not influenced by BtuCD. For smFRET-Q experiments, in addition to the appended C-terminus cysteine and His_6_-tag, a (Gly_4_Ser)_2_ sequence was added and served as a spacer to allow maximum mobility of the surface tethered BtuF*. This was followed by an AviTag (Avidity LLC, Aurora, CO) for covalent biotinylation for surface immobilization. For two-color ensemble FRET experiment, cystine-less BtuCD was created by replacing all native cysteine residues with serine for site-specific labeling34. Periplasmic residue Q111 was subsequently mutated to cysteine for fluorophore labeling. For all other experiments, wild type BtuCD was used. Tandem repeat BtuC-BtuC-BtuD_2_ constructs (EE, EQ and QQ)16 were made by gene synthesis (Genscript, Piscataway, NJ), adding a (Gly_4_Ser)_5_ linker at the C terminus of BtuD, followed by another BtuD sequence. EE contains two wild type BtuD copies; EQ contains one wild type BtuD followed by another BtuD with E159Q mutation; QQ contains two BtuD E159Q copies.

### Protein expression, purification.

Following induction at OD_600_=0.6 and overnight expression at 25 °C in *E. coli* C43 (DE3) cells (EMD Millipore, Novagen), cells were collected by centrifugation. All BtuF constructs were purified directly from whole cell lysate (obtained by sonication) via Ni-NTA chromatography followed by size-exclusion chromatography (Superdex 200 10/300 GL, GE Healthcare) in final buffers of 20-50 mM Tris-HCl, 100-150 mM NaCl, 0-1 mM Tris(2-carboxyethyl)phosphine (TCEP) at pH ∼7.4. All BtuCD constructs were purified from membrane pellets in 0.1% Lauryldimethylamine-oxide (LDAO) using the same purification techniques as above. Membrane scaffold protein (MSP) construct 1E3D1 that was used for nanodisc reconstitution was purified essentially as previously described50.

### Nanodisc and proteoliposome reconstitutions

Nanodisc-BtuCD was used in all experiments except for the liposome transport assay (Fig. S2e-f). For nanodisc reconstitution, *E. coli* polar lipid extract (Avanti) was dried and rehydrated to 40 mg/mL (∼50 mM) in 100 mM NaCl, 1 mM TCEP, 100 mM sodium cholate and 20 mM Tris-HCl at pH 7.4 and mixed with MSP1E3D1 and BtuCD. Final concentrations were: 5 mM lipid, 12 µM BtuCD, 72 µM MSP1E3D1, and 30mM sodium cholate. To minimize aggregation, 4% glycerol was also added. The mixture was incubated at 4 °C for 1 hour with gentle agitation. 400 mg of wet Bio-Beads SM-2 (BioRad) per 500 μL of mixture was added and the mixture continued to incubate overnight to remove detergents. Bio-beads were subsequently removed. Reconstituted nanodisc-BtuCD was purified from Ni-NTA agarose and then size-exclusion chromatography on a Superdex 200 column.

For liposome reconstitution, liposomes were created from a 3:1 mix of E. coli polar lipid extract and 1,2-Dioleoyl-sn-glycero-3-phosphocholine (DOPC), reconstituted with membrane proteins, and loaded with BtuF and B_12_ as previously described51. Concentrations of 10 μM unlabeled BtuF and 25 μM B_12_ were used in the loading step.

### Fluorophore labeling and biotinylation

BtuF and niBtuF were labeled with Alexa Fluor 546 (AF546) C_5_ Maleimide (ThermoFisher) according to the manufacturer’s manual. BtuCD Q111C was used in two-color FRET ensemble experiments and was labeled with Alexa Fluor 647 (AF647) C_5_ Maleimide prior to nanodisc reconstitution. For smFRET experiments, BtuF with AviTag is biotinylated according to the protocol offered by Avidity with minor changes. Biotinylated BtuF was then labeled with AF 546. All labeled proteins were purified from Ni-NTA agarose followed by size-exclusion chromatography to eliminate free fluorophores and aggregates.

### ATPase activity measurements

ATPase activity was measured by using Malachite Green Phosphate Assay Kit (Sigma-Aldrich). ATPase activity of BtuCD in nanodiscs was carried out in a 500 μL reaction buffer containing 50 mM Tris-HCl, pH 7.5, 150 mM NaCl, 2 mM Mgcl_2_ (Cacl_2_ in experiment testing Ca-ATP hydrolysis). Then, 50 nM nanodisc BtuCD was mixed with 1 µM BtuF and 5 µM B_12_ in reaction buffer at 25 °C. ATP was added at t = 0 to initiate the reaction. Once ATP was added, 80 μL aliquots were removed at various time points (1, 2, 3, 4, 5 and 6 min) and mixed with 20 μl working reagent containing malachite green in a 96-well plate to quench the hydrolysis reaction. Samples were then incubated for 30 min for color development. The amount of inorganic phosphate (Pi) at each time point was determined by taking the absorbance at 620 nm using a plate reader (BioTek Synergy Neo 2) and comparing to a standard Pi concentration plot.

### Ensemble FRET experiments in bulk solution

Fluorescence measurements in bulk solution (1.1 mL cuvette) were made on a PC1 photon-counting spectrofluorimeter (ISS Inc., Champaign, IL), using an excitation wavelength of 540 nm and an emission wavelength of 573 nm to measure the intensity of AF546 on BtuF*. For single-color FRET experiments that use AF546-labeled BtuF and unlabeled BtuCD, total fluorescence was calculated from the formula *I_Total_*=*I_VV_*+2*G*\**I_VH_* and anisotropy was calculated from the formula *r*=(*I_VV_*−*G*\**I_VH_*)/*I_Total_*, where *IVV* is the fluorescence intensity recorded with excitation and emission polarization both in the vertical position, *I_VH_* the fluorescence intensity with the emission polarization aligned in the horizontal position, and *G*=*I_HV_*/*I_HH_*;. For two-color FRET experiments that used AF546-labeled BtuF and AF647-labeled BtuCD, additional fluorescent emission at 665nm was measured simultaneously.

### Estimation of the AF546-B_12_ Förster distance and travel distance of B_12_ during transport process

The estimate of the AF546-B_12_ Förster distance *R*_0_ used to generate Fig. 1b was calculated from the observed quenching level (56% quenching) of BtuF* at saturating B_12_ concentration using the following equation:

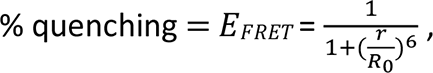

where *E*_*FRET*_ is the FRET efficiency (which is also the % quenching), and *r* the AF546-B_12_ distance. The calculated Förster radius, R_0_, was then used to plot the theoretical AF546 fluorescent intensity as a function of the transit distance, which taken to be the projection of the AF546-B_12_ distance along the axis running through the BtuCD-F complex (PDB 4FI3) permeation pathway (Fig. 1b).

### smFRET-Q imaging using Total Internal Reflection Fluorescence (TIRF) microscopy

Biotinylated-BtuF* described in previous section was tethered to the surface of a quartz microfluidic flowcell that was pre-functionalized with polyethylene glycol (PEG)/biotin-PEG (mPEG-SVA MW3400 and Biotin-PEG-SVA MW5000, respectively, Laysan Bio) and pre-derivatized with streptavidin (ThermoFisher) *via* a streptavidin-biotin-streptavidin bridge^52,53^. In this tethering process, 100 pM biotinylated-BtuF* was incubated in the flowcell for 5 minutes at room temperature. Untethered BtuF* was removed by washing the flowcell with Sample Buffer (SB), composed of 50 mM Tris, 150 mM NaCl, 2 mM Mgcl_2_ (or Cacl_2_ for experiments performed using Ca-ATP), 50 μM EDTA, pH = 7.4.

Just prior to imaging, flowcells containing tethered, biotinylated BtuF* were washed with Imaging Buffer 1 (IB1), composed of the same components as SB, but with the addition of: (*i*) nucleotides, when necessary (MgATP, CaATP, MgAMPPNP, or minus nucleotide as listed in **Supplementary Information Table 1**); (*ii*) an oxygen-scavenging system (2.5 mM Protocatechuic acid (PCA, pH ∼8, Sigma-Aldrich), 60 nM Protocatechuate-3,4-dioxygenase (PCD, Sigma-Aldrich))^54^; (*iii*) a triplet-state quencher cocktail (1 mM cyclooctatetraene (COT) and 1mM 4-nitrobenzyl alcohol (NBA))^55^; and (*iv*) a reducing and oxidizing system (2.5 mM ascorbic acid (neutral pH) and 2.5 mM methyl viologen)^56^.

smFRET-Q imaging was conducted using a wide-field, prism-type, TIRF microscope. The AF546 fluorophores attached to flowcell-surface-tethered, biotinylated BtuF* molecules in the field-of-view (FOV) were excited with a 30 mW beam from a continuous-wave 532 nm laser (Gem 532; Laser Quantum). Fluorescence emission from each AF546 was collected through a 60*×*, 1.2 NA water immersion objective (Nikon) on an inverted microscope stand (TE2000; Nikon), and imaged on a 1024 *×* 1024 pixel, back-illuminated electron-multiplying charge-coupled device (EMCCD) camera (iXon Ultra 888; Andor) using 2*×* pixel binning. Movies were collected at a time resolution of 60 msec per frame with an estimated ∼0.1 kW/cm^2^ illumination power density at the sample interface.

Initially, the fluorescence intensity from thousands of surface-tethered, biotinylated BtuF* molecules in the FOV in IB1 were recorded for 30 frames (1.8 sec). The laser beam was then shuttered and the recording was turned off. Shortly thereafter, Imaging Buffer 2 (IB2), composed of the same components as IB1, but with the addition of the specified concentrations of B_12_ and BtuCD (as listed in **Supplementary Information Table 1**) was stopped-flow delivered into the flowcell with the laser beam still off. Stopped-flow delivery took place 30 seconds after BtuCD was added to the B_12_- and ATP-containing IB2, which initiated the ATPase reaction. IB2 was subsequently incubated in the flowcell for 60 seconds while the laser beam remained off. The laser was then turned on again and recording of the same FOV continued for 1000 frames (1 min). Triplicate experiments were performed for each experimental condition (listed in **Supplementary Information Table 1**) using different flowcells on different days.

### smFRET-Q data analysis

Because vitamin B_12_ is a non-photon-emitting FRET acceptor, we used changes in the fluorescence intensity of the AF546 FRET donor fluorophore as a proxy reporter for changes in the efficiency of FRET (E_FRET_). We therefore identified the fluorescence intensity originating from individual AF546 fluorophores attached to individual, surface-tethered, biotinylated BtuF* molecules in each TIRF movie by identifying the local maxima in an image of the single-frame delay autocorrelation function of each pixel, with the equation

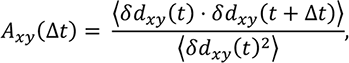

where *d_xy_*(*t*) is the camera intensity value of the pixel at (x,y) at time t, ⟨*X*(*t*)⟩ is the time average of *X*(*t*), δ*X*(*t*) = *X*(*t*) − ⟨*X*(*t*)⟩, and, in this case, the time delay was Δ*t* = 1 frame. This autocorrelation image approach has the benefit of removing intensity noise from the search process that is noise from the search process that *vs.* time trajectories were generated by estimating the fluorescence intensity above the background intensity in each frame for each identified fluorophore using an iterative, maximum likelihood formula, 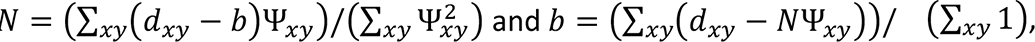, where *xy* denotes a pixel, *N* is intensity, *b* is background, *db* is the value of the image data, Ψ is an isotropic 2D Gaussian-shaped point spread function, and the sums are performed over the 7x7 pixel local region of each fluorophore. In order to be able to directly compare trajectories obtained from different surface-tethered, biotinylated BtuF* molecules, we normalized the AF546 fluorescence intensity of each 1000-frame trajectory (recorded using BtuF* in IB2). We accomplished this by using the initial 30 frames of AF546 fluorescence intensity *vs.* time trajectory (recorded using *apo* BtuF* in IB1) we recorded for each surface-tethered, biotinylated BtuF* molecule to calculate the average AF546 fluorescence intensity for each trajectory. Subsequently, we normalized the AF546 fluorescence intensity recorded in each frame of the 1000-frame trajectory (recorded using BtuF* in IB2) corresponding to the same BtuF* molecule by the average AF546 fluorescence intensity obtained for that trajectory. Each resulting, 1000-frame, normalized AF546 intensity vs. time trajectory represents the fractional intensity of the AF546 FRET donor fluorophore that is quenched *via* FRET to the vitamin B_12_ FRET acceptor as a function of time and therefore acts as a proxy reporter for the corresponding E_FRET_ *vs.* time trajectory, as described in the main text of this article. The initial 30 frames of AF546 fluorescence intensity *vs.* time trajectory (recorded *apo* BtuF* signal in IB1) as well as any frames following AF546 photobleaching were removed before further processing of the trajectories (as shown in **Fig. 1f-h**).

Only fluorescence intensity *vs.* time trajectories that satisfied the following criteria were kept for further analysis: (*i*) a signal-to-background ratio, computed as the difference in average fluorescence intensity before and after photobleaching divided by the standard deviation of the fluorescence intensity after photobleaching57, of greater than 3; (*ii*) one instance of 54 +/-5% quenching at minimally two time points, which distinguished functional BtuF* molecules from non-functional BtuF* molecules and any contaminants emitting or scattering light within the wavelength range of our emission filter (575 ± 40 nm) (typically less than 10% of the trajectories from any experimental condition were removed due to this criterion); (*iii*) the absence of significant photophysical noise, such as fluorophore blinking, as identified by eye; and (*iv*) only a minimal amount of defocusing induced by the stopped-flow delivery procedure. Non-functional BtuF* molecules mentioned in (*ii*) can possibly be due to preferred orientation, misfolding, degradation at N-terminal end, or aggregation that prevented B_12_ binding. The level of defocusing mentioned in (*iii*) for each trajectory was quantified by the level of maximum intensity. Trajectories with maximum normalized intensities that were greater than 1.1 and lasted for more than one continuous frame were not included.

The normalized, AF546 fluorescence intensity *vs.* time trajectories from each movie were analyzed together using a variational Bayesian Hidden Markov Model (HMM) to yield a global model of the single-molecule observations and dynamics present in each replicate movie of each experiment58,59 Two-state HMMs were used to analyze the B_12_ binding data in **Fig. S5a** and the resulting transition matrices were used estimate the values of *k_binding_* and *k_release_*. For the experiments listed in **Supplementary Information Table 1**, we initially analyzed all datasets using global HMMs with two through six states. Visual inspection of the results of these analyses showed that a minimum of four states was required to adequately describe experiments conducted with 1-2 µM BtuCD (WT and EE constructs) and 0.5 mM Mg-ATP, regardless of the concentration of B_12_ (**Supplementary Information Table 1**). However, a minimum of five to six states became necessary to adequately describe trajectories from experiments using reduced BtuCD and Mg-ATP concentration or experiments using EQ and QQ constructs. For trajectories from experiments conducted with Mg-AMPPNP, Ca-ATP and no nucleotide, six states no longer adequately described the states with dequenched intensity levels. While much of the signal complexity that necessitates the use of global HMMs with several states is due to non-uniform broadening of fluorescence intensity levels in this smFRET-Q study, we found that for all experiments (except those conducted with Mg-AMPPNP, Ca-ATP and no nucleotide) analyzed using four-through six-state global HMMs, two significantly dequenched states were always found near 0.82 and 1.0, while the rest of the states were tightly clustered near 0.46. Because of this effect was observed independent of the molecular details of BtuCD, the states near 0.46 were clustered into a common state, with a mean centered at the average of the individual intensity state means weighted by their fractional occupancies, in order to simplify the model and to focus exclusively on the transport process. The clustered, low intensity state was assigned to the L state, and the two dequenched states were assigned to the I and H states as described in the main text. The mean normalized intensities of the L, I, and H states for each replicate in each experimental condition are given in **Supplementary Information Figure 11**.

To quantify the kinetics of the transitions between these three states, a three-state global HMM was estimated where the intensity state values were fixed to the mean of the normalized intensity values determined for each replicate and for experimental condition for the L, I, and H states. This analysis gave idealized state versus time trajectories (i.e., Viterbi paths) that were used to generate 2D surface contour surface plots of the time evolution of population FRET and transition density plots. For these plots, triplicate datasets for each experimental condition were first combined, and then analyzed using the HMM modeling processes described above. For the surface contour plots showing transitions into the H state, the first data point of each sampling of the H state from all of the trajectories were superimposed. For the surface contour plots showing transitions out of the I state, the last data point of each sampling of the I state from all of the trajectories were superimposed. The surface contour plots show either only one state or all states (left and right panels, respectively, in Fig. 2a-b) before and after the specified state, as indicated in figure legends. Surface contours are colored as denoted in the population color bars. For transition density plots, all inter-state transition events identified in the idealized state versus time trajectories for all of the replicate datasets recorded under an experimental condition were superimposed and plotted as the normalized AF546 fluorescence intensity before the transition versus the normalized AF546 fluorescence intensity after the transition.

Within these three-state global HMMs, the kinetics of transitions between the L, I, and H states are quantified with a 3 x 3 transition probability matrix. Th transition probabilities, *p*, within that matrix were converted to the rate constants, *k*, shown in Fig. 3 using the equation

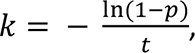

where *t* is the time interval between data points (60 ms). Error bars were standard deviations calculated from corresponding rate constants determined from triplicate experiments. For experiments using Mg-AMPPNP, Ca-ATP, and no nucleotide, in which dequenching events were rare, the means of the L, I, and H states were first obtained from the combined triplicate datasets obtained from the reference experiment (the first row in Supplementary Information Table 1), and then fixed while estimating a three-state, global HMM to obtain the 3 x 3 transition probability matrix for the L, I, and H states, and then convert those transition probabilities into the corresponding rate constants.

To assess the extent of the variation we observed in the means of the normalized intensity values for the L, I, and H states within and across experimental conditions, we plotted the means of the normalized intensity values for the L, I and H states (Fig. S11). For the Mg-AMPPNP, Ca-ATP, or no nucleotide experimental conditions, only trajectories that exhibited dequenching events were included in the state estimation in this plot. The majority of the means of the normalized intensity values for the L, I, and H states are between 0.42 and 0.48, 0.78 and 0.86, and 0.96 and 1.07, respectively. This plot demonstrates that the means of the normalized intensity values for the L, I, and H states are well-separated from each other, allowing the L, I, and H states to be clearly distinguished and assigned. It should be noted that the means of the normalized intensity values for the I state in the Mg-AMPPNP, Ca-ATP, or no nucleotide experimental conditions, while still within the 0.78 – 0.86 range, were systematically lower than in the other experimental conditions. This systematic decrease in the means of the normalized intensity values for the I state was accompanied by faster rate constants for exiting the I state in these experimental conditions *versus* the reference experimental conditions (Fig. 3). Together, these observations suggest that the I state in the Mg-AMPPNP, Ca-ATP, or no nucleotide experimental conditions (which we refer to as the I’ state) represents a potentially different transport intermediate than the I state that we observe in the other experimental conditions, as discussed in the main text of this article.

## Supplementary Information

**Fig. S1.**
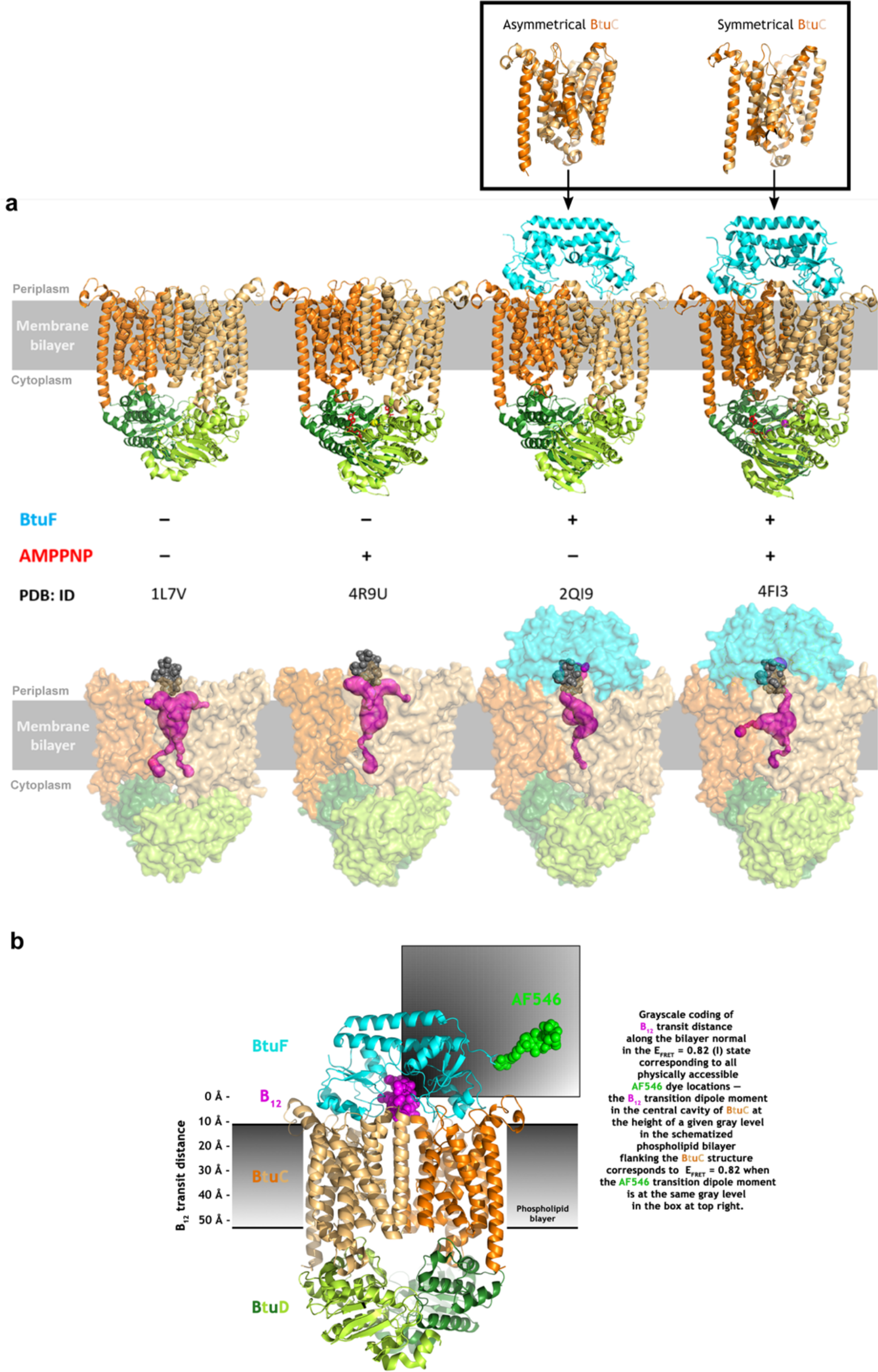
BtuCD internal cavity properties in available crystal structures and systematic calibration of B_12_ transit distance during the L -> I transition. **a**, Crystal structures of BtuCD/BtuCD-F complex either in *apo* form or bound by AMPPNP shown in ribbon diagrams (upper) and surface representations (lower). In the lower panel, B_12_ molecule is shown in grey space-filling representation and is aligned with entrance pathway of BtuC dimer. The magenta spheres show the internal cavities in BtuC dimer in each structure and all channels connecting that cavity to either the cytoplasmic or periplasmic surface of BtuCD-F, as analyzed using the Caver algorithm. b, The fluorescence intensity change during the L -> I transition corresponds to B_12_ movement approximately into the middle of the membrane for all physically reasonable positions of the AF546 fluorophore. The grayscale color in the schematized phospholipid bilayer flanking the ribbon diagram of BtuCD-F* encodes the location of B_12_ within the central cavity of the BtuC dimer if the transition dipole moment (TDM) of AF546 is located at a position with the same grayscale level in the rectangular box at the upper right. The transit distance was calculated from the molecular coordinates using the Förster equation for FRET efficiency assuming that, in the L state, B_12_ is located at its crystallographically observed binding site on BtuF, while in the I state, it moves directly along the approximate 2-fold symmetry axis at the center of BtuCD, which is parallel to the membrane normal (*i.e.*, the axis running directly through the membrane perpendicular to its surfaces). For any assumed location of the TDM of AF546 in this molecular coordinate system, the observed 0.54% quenching of AF546 in the L state was used to calculate the corresponding value of the Förster radius, R_0_, which was in turn used to calculate the location of B_12_ corresponding to the 0.18% quenching observed in the I state. The FRET efficiency calculation assumes random orientation of the TDM of AF546 relative to that of B_12_, which produces cylindrical symmetry around the approximate 2-fold axis at center of the BtuCD assembly. Therefore, the calculation covers all physically reasonable locations of the TDM of AF546 even though the schematic diagram only shows them in a single plane parallel to the membrane normal. The ribbon diagram shows the crystal structure of BtuCD-F in the absence of B_12_ (PDB id 4DBL) with BtuC colored shades of orange, BtuD colored shades of green, and BtuF colored light blue. B_12_, shown in magenta space-filling representation in its binding site in BtuF, was positioned based on least-squares alignment of BtuF in a crystal structure of the BtuF•B_12_ complex in the absence of BtuCD because it has not been possible to visualize B_12_ in any crystal structure containing BtuCD.

**Fig. S2.**
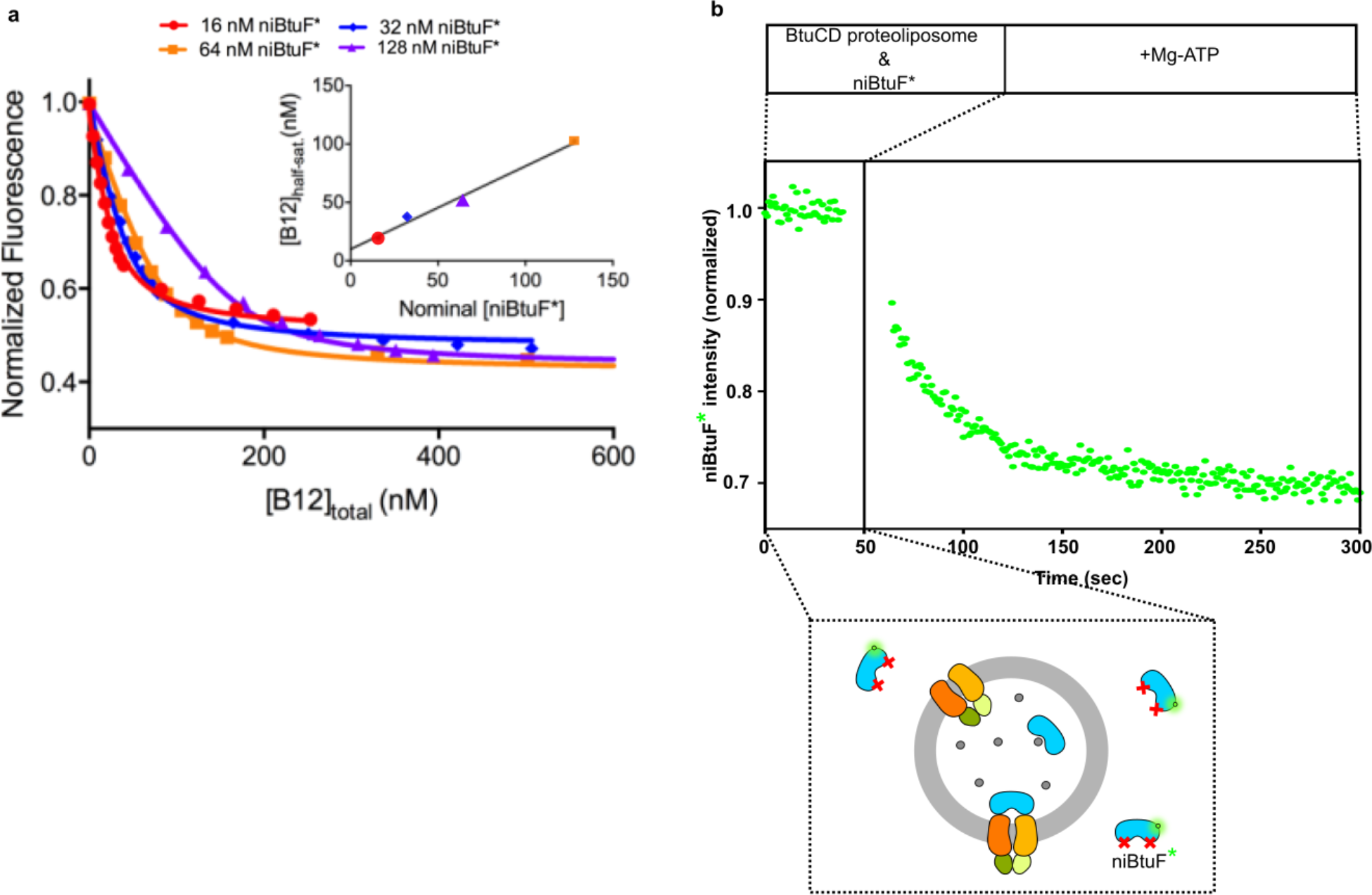
ATP-dependent extrusion of B_12_ by WT BtuCD reconstituted into phospholipid vesicles. **a**, Characterization of the B_12_ binding affinity of “<u>n</u>on-interacting” BtuF* (niBtuF* — AF545-labelled E72A-E202A-BtuF*), a double mutant that shows no significant interaction with BtuCD34 under relevant experimental conditions and can therefore serve as a fluorescence biosensor for B_12_ in transport experiments. The graph shows titrations of B_12_ onto four different concentrations of niBtuF* monitored by the quenching of AF546 fluorescence, which demonstrate that niBtuF* has very similar B_12_ binding affinity and quenching properties as WT BtuF*. The *K_d_* for B_12_ binding by niBtuF* estimated from global non-linear least-squares curve fitting of these data is 13 ± 1 nM. The inset shows B_12_ concentration at half-saturation inferred from non-linear curve-fitting of individual titrations *versus* nominal BtuF* concentration in the different titrations, which yields a stoichiometry estimate of 1.4 ± 0.1 niBtuF* and a *K_d_* estimate of 10 ± 5 nM based on linear least-squares curve-fitting to the equation [B_12_]_50%_ saturation = *s*-1(*K_d_* + ([BtuF*_total_]/2)). **b**, Demonstration of efficient ATP-dependent B_12_ transport by WT BtuCD reconstituted into phospholipid vesicles. The solution contained 40 nM niBtuF* in the external buffer and 300 nM BtuCD in proteoliposomes loaded with 25 µM B_12_ and 10 µM BtuF, as schematized below the graph. Addition of 0.5 mM Mg-ATP at t = 50 sec leads to efficient extrusion of B_12_ from the proteoliposomes as monitored by quenching of the external niBtuF*. All experiments were conducted at room temperature.

**Fig. S3.**
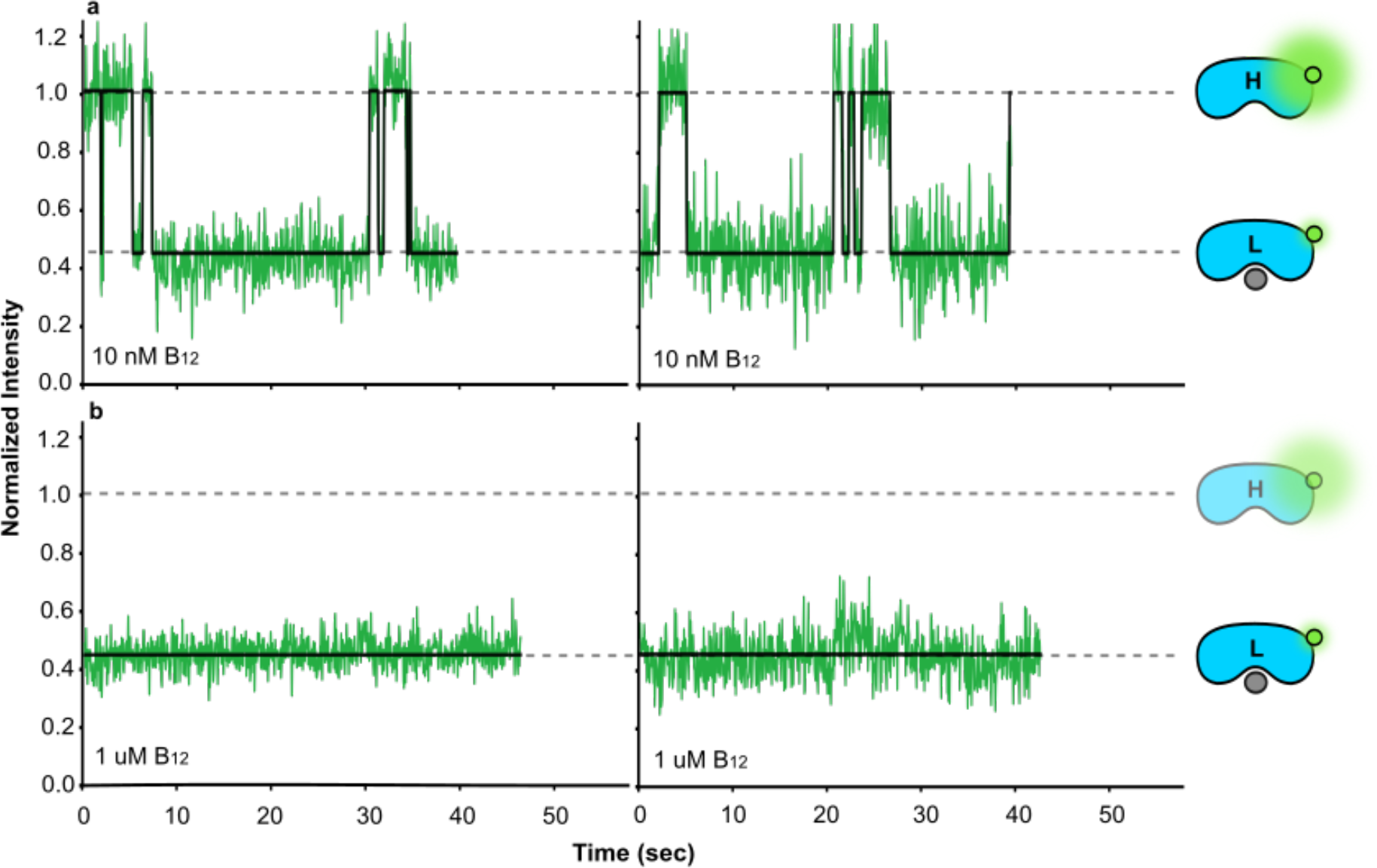
Characterization of B_12_ binding kinetics by smFRET. Representative trajectories showing BtuF* intensities in the presence of 10 nM (**a**) or 1 µM (**b**) B_12_. Kinetic parameters were derived from HMM analyses of BtuF* trajectories in the presence of 10 nM B_12_, yielding *k_on_* = 0.065 ± 0.005 s-1●nM-1 and *k_off_* = 0.130 ± 0.01 s-1. The *K_d_* calculated using these rate constants is 2 nM.

**Fig. S4.**
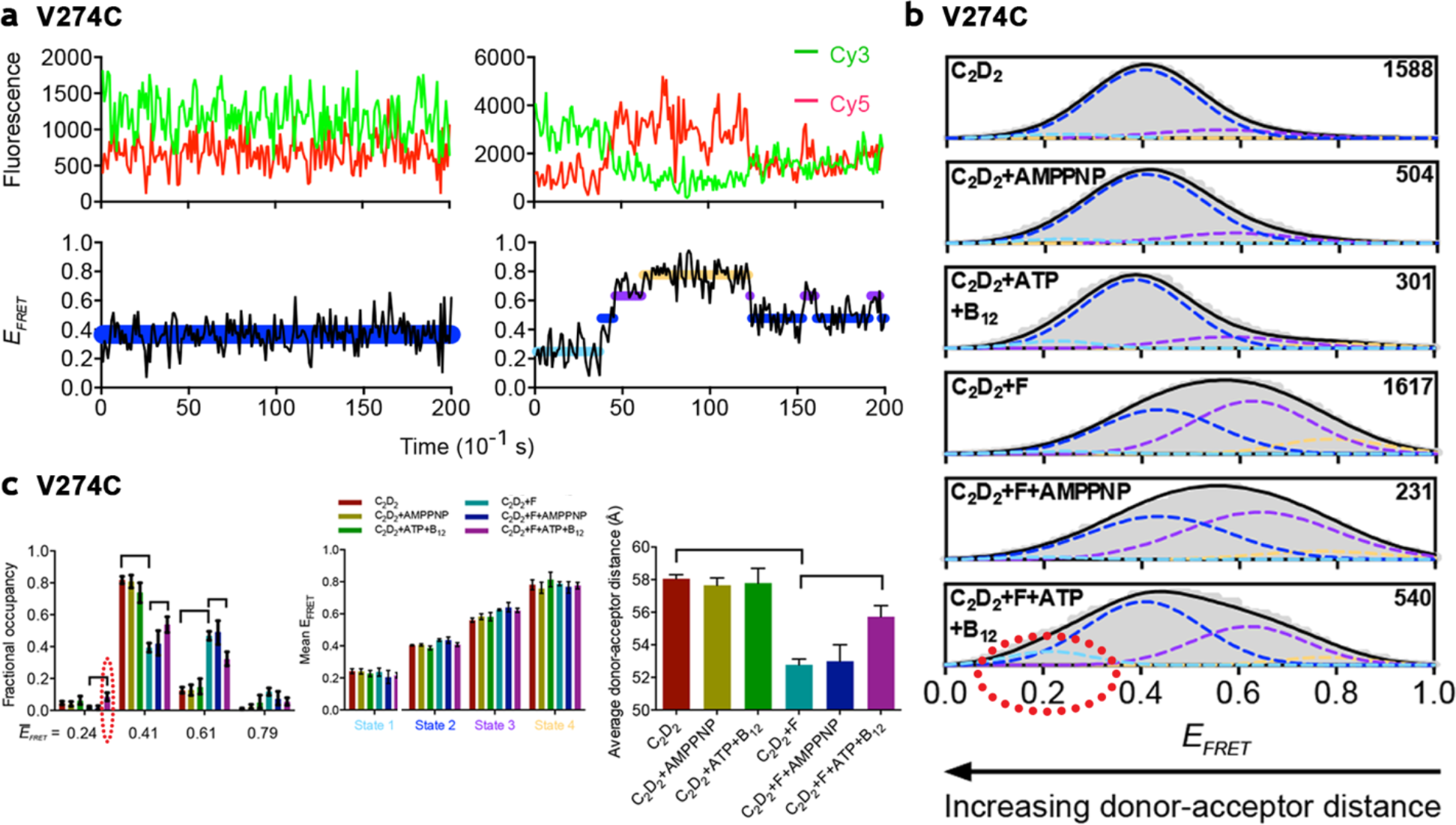
Dual-color smFRET experiments on BtuCD-F complexes demonstrate that ATP binding does not induce a significant conformational change whereas ATP hydrolysis induces transient occupancy of a conformational state with a wider opening of the BtuC domains. The V274C mutation was introduced into a cysteine-reduced variant of BtuCD34 in which the single remaining native cysteine is buried (C279 in BtuC). The protein complex was purified in LDAO, stochastically labeled at the V274C site with a 50:50 mixture of Cy3 and Cy5 maleimides, and repurified to remove unreacted fluorophores. The labeled complex was then reconstituted into nanodiscs formed from *E. coli* polar lipids and the MSP1E3D1-T277C scaffold peptide34 after covalently derivatizing it with biotin maleimide. The BtuCD-containing nanodiscs were then surface-tethered for collection of two-color smFRET data using equivalent methods^60^ to those employed for our smFRET-Q experiments. The two-color smFRET data collected from BtuCD labeled with Cy3/Cy5 at the V274C site in BtuC demonstrate that addition of AMPPNP either in the absence or presence of BtuF produces minimal changes in the inter-fluorophore distance distribution, while addition of ATP together with B_12_ in the presence of BtuF produces ∼6% occupancy of a state with mean E_FRET_ = 0.24. This value corresponds to an ∼9 Å wider opening of the BtuC dimer compared to the widest state with substantial occupancy under all other conditions (assuming a distance of half-maximal E_FRET_ of 55 Å for Cy3/Cy5). This state, which is highlighted by dotted red ellipses in panels b-c, also shows evidence of enhanced occupancy in in the presence of ATP in the absence of BtuF but at a lower level than in the presence of both ATP and BtuF. The molecular geometry is illustrated in the ribbon diagrams in Fig. S5a below. a, Baseline- and crosstalk-corrected FRET donor (Cy3) and acceptor (Cy5) fluorescence intensity (arbitrary units) *versus* time (sec) trajectories (top row) and the corresponding E_FRET_ *versus* time trajectories (bottom row) from BtuCD complexes stochastically labeled with Cy3/Cy5 at the V274C site in BtuC. The trajectories on the left photobleach prior to exhibiting any transitions, while those on the right exhibit transitions prior to photobleaching. The idealized E_FRET_ *versus* time trajectories (*i.e.*, Viterbi paths) determined from a 4-state HMM^60^ are superimposed over the experimentally determined E_FRET_ *versus* time trajectories, with different colors used to represent the different inferred E_FRET_ states. The procedures used to select the 4-state HMM and estimate the related parameters are described below in this legend. **b,** E_FRET_ histograms (normalized to the maximum value) for trajectories recorded under different experimental conditions from samples stochastically labeled with Cy3/Cy5 at the V274C site in BtuC, with the number of trajectories recorded for each condition shown at the top right corner of each histogram. Population distributions for the individual states of the 4-state HMMs60 (colored dashed lines) and their population weighted sum (solid black line) are superimposed. Each HMM state is represented by a Gaussian centered at the mean of all E_FRET_ values assigned to that state, with its standard deviation set equal to the standard deviation of all E_FRET_ values assigned to that state and its amplitude scaled to the average fractional occupancy of that state. Note that the E_FRET_ distribution does not change in the datasets collected in the presence of Mg•AMPPNP compared to the datasets from the same complexes collected in the absence of nucleotide, suggesting that ATP binding induces little to no changes in the conformational states of BtuCD or BtuCD-F. In contrast, there is a significant increase in the fractional occupancy of the conformational state corresponding to E_FRET_ ≈ 0.24 in the presence of both ATP and BtuF (dotted red ellipses in the E_FRET_ histogram in panel **b** and the bar graph of fractional occupancies in panel **c**), which is consistent with ATP hydrolysis inducing occupancy of a conformational state with a significantly wider opening of the TMDs. The relative transition probabilities in the 4-state HMMs estimated for these datasets show that transitions occur predominantly between neighboring E_FRET_ states (data not shown), meaning those centered at the closest E_FRET_ values; this pattern is consistent with movement between stereochemically adjacent conformational states with progressively narrower/wider separation between the TMDs. **c,** The fractional occupancies (left), mean E_FRET_ values (center), and corresponding estimated donor-acceptor distances (right) for each of the four modeled E_FRET_ states under each experimental condition for the fluorophores at the V274C site in BtuC. The E_FRET_ states are numbered from 1 to 4 in increasing order of their average E_FRET_ values, so that a state with a larger number corresponds to a shorter donor-acceptor distance. The error bars represent 95% confidence intervals obtained by bootstrapping, while the brackets highlight statistically significant differences. The 4-state HMM used to analyze the data described above was chosen based on parallel analyses of the datasets collected under the different experimental conditions. In each individual analysis, the ebFRET software suite60 was used to model the full set of trajectories collected under a single experimental condition. This program employs an empirical Bayesian framework to estimate a single HMM that simultaneously describes all of the E_FRET_ *vs.* time trajectories in the dataset being analyzed, and it thereby assigns all of the measured E_FRET_ values to a set of common states corresponding to different separations between the donor and acceptor fluorophores. The complete set of trajectories from each individual experimental condition was systematically analyzed using five different HMM models ranging from two states to six states to determine which model maximizes the evidence per trajectory, which is thus the most parsimonious description of that dataset. This set of analyses led to the conclusion that the 4-state model most often provides the most parsimonious description of the totality of the data collected under the various experimental conditions for BtuCD stochastically labeled with Cy3 and Cy5 at the V274C site in BtuC. Therefore, the 4-state HMM was selected for modeling of the datasets collected under all experimental conditions. Equivalent analyses performed on data collected from BtuCD labeled with the same fluorophores at each of four additional sites (**Figs. S5-S6** below) similarly support the conclusion that a 4-state HMM provides the most parsimonious description of the totality of the smFRET data collected from all labeling sites.

**Fig. S5.**
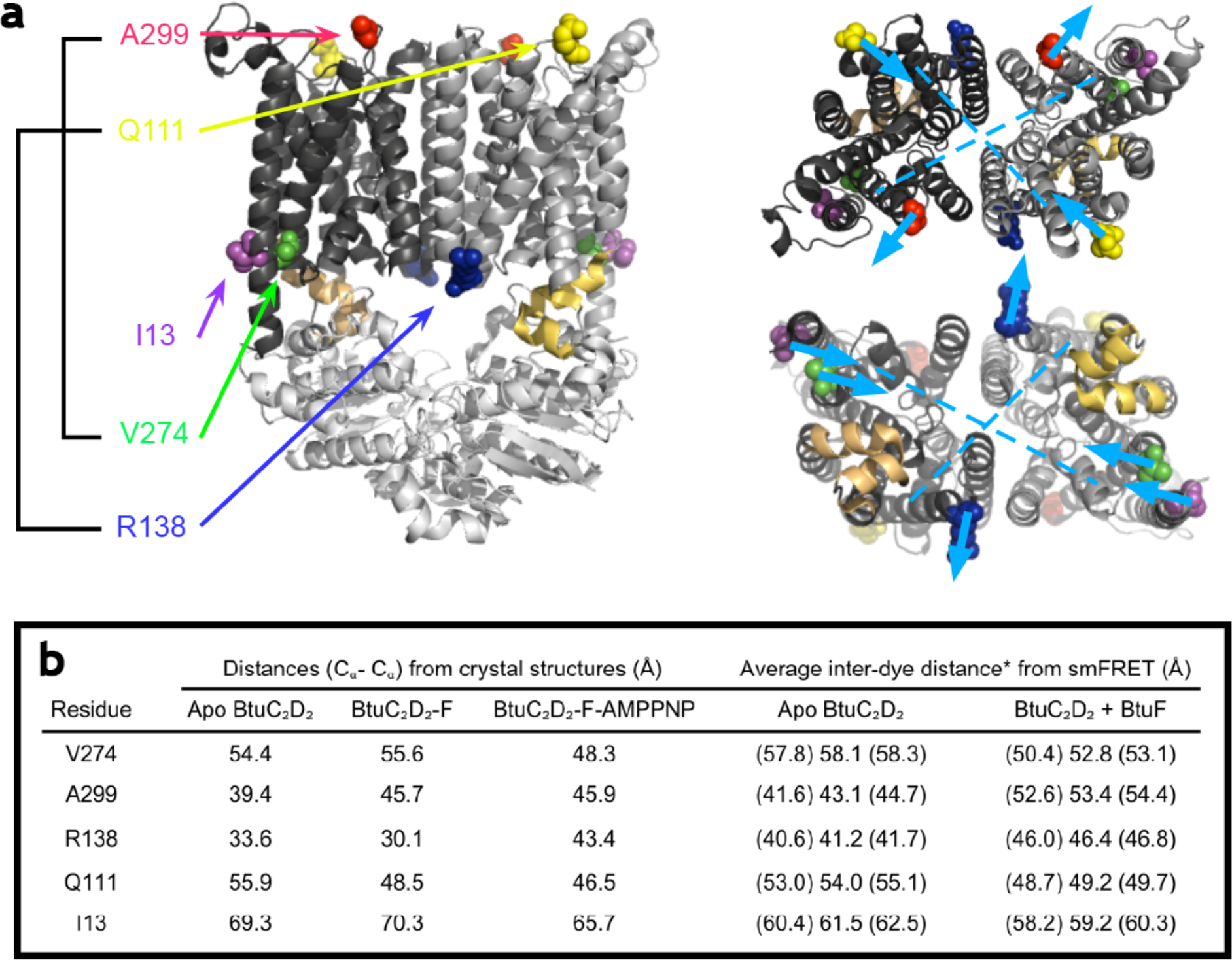
Changes in average inter-fluorophore distance upon BtuF binding. **a**, The ribbon diagram on the left illustrates the fluorophore labeling sites used in the dual-color smFRET experiments shown in **Figs. S4** and **S6,** while the ribbon diagrams on the right schematize the direction of the distance changes at those site upon BtuF binding, calculated from mean E_FRET_ values. BtuCD is viewed from the plane of the plasma membrane (left), from the periplasm (top right), or from the cytoplasm (bottom right), with the residues mutated to cysteine for fluorophore labeling shown as colored spheres and the NBD-interacting helices in BtuC shown in beige. V274 and A299 are located on TM helix 9, while R138 and Q111 are located on TM helix 4, and I13 is located on TM helix 1. The light blue dashed lines show reference vectors between the centers of TM helices 9 and 4, while the light blue arrows schematize the directions of the inferred movements of the labeled residues in response to BtuF binding. **b**, Table comparing inter-residue distances calculated from available crystal structures to mean donor-acceptor distances calculated from smFRET experiments conducted on each labeled construct. The numbers in parentheses correspond to 95% confidence limits determined using bootstrapping. Donor-acceptor distances were calculated from mean E_FRET_ values using the Förster equation assuming an R_0_ of 55 Å. The *apo* BtuCD, BtuCD-F, and BtuCD-F•AMPPNP structures used for the calculations have PDB ids 1L7V, 2QI9, and 4FI3, respectively.

**Fig. S6.**
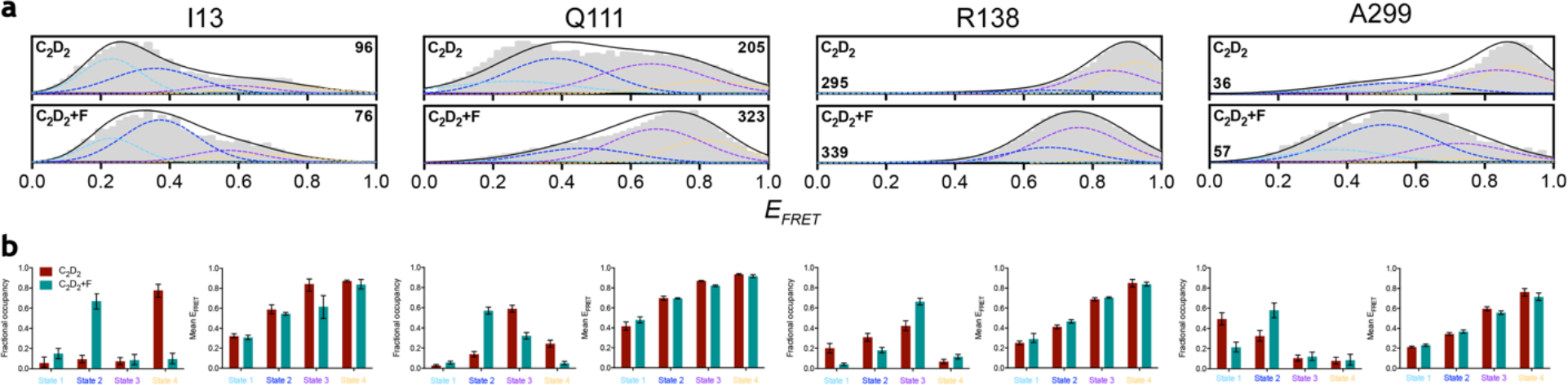
E_FRET_ histograms and state-occupancy statistics for additional dual-color smFRET experiments. Data are shown for all of the labeling sites presented in **Fig. S5** except for V274C (which is shown in **Fig. S4**). The data were generated and analyzed using 4-state HMMs as described in **Fig. S4** except that the BtuCD construct used to introduce the single-site cysteine mutations had all native cysteine residues removed including C27934. **a**, E_FRET_ histograms for the dynamic trajectories recorded under different experimental conditions, with the number of trajectories recorded for each condition shown at the top right or bottom left corner of each histogram. **b,** Fractional occupancies (left) and mean E_FRET_ values (right) from the 4-state HMM estimated for the dataset above each pair of graphs. E_FRET_ (conformational) states for each donor-acceptor pair are again numbered from 1 to 4 in increasing order of their average E_FRET_ values, so that a state with a larger number corresponds to a shorter donor-acceptor distance. The error bars represent 95% confidence intervals obtained by bootstrapping.

**Fig. S7.**
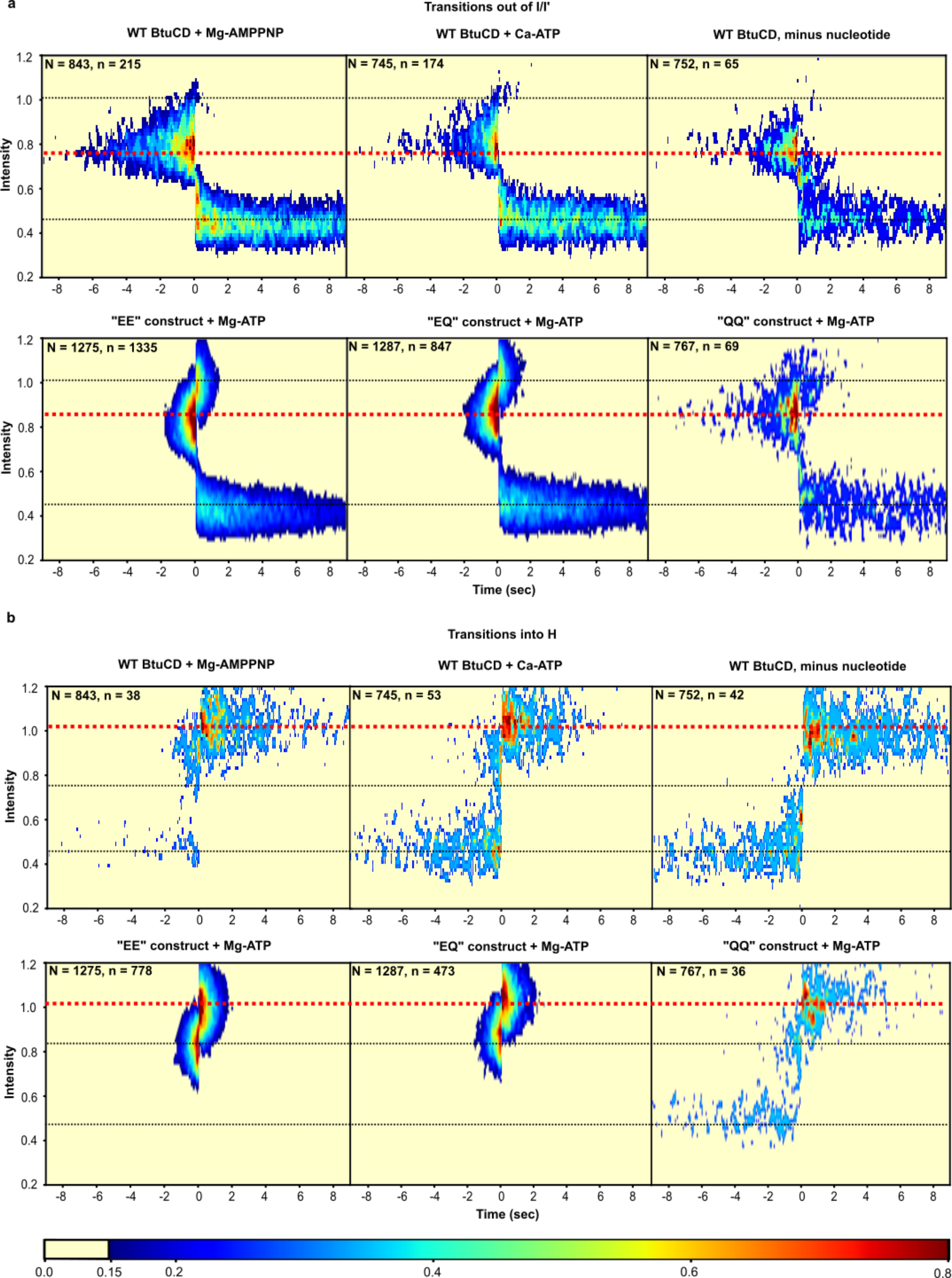
ATP hydrolysis drives unidirectional conformational transitions. Synchronized 2D contour plots equivalent to those shown in Fig. 2a**-b** in the main text but for experiments conducted with WT BtuCD in the absence of nucleotide or the presence of nonhydrolyzable analogs (top plots in each section) or using tandem BtuCD constructs instead of WT BtuCD in the presence of ATP (bottom plots in each section). The EE, EQ, and QQ tandem BtuCD constructs are schematized and characterized in **Fig. S10** below. Transitions out of the I state or I’ states are shown in part **a**, while transitions into the H state are shown in an equivalent layout in part **b**. The very small number of transitions into the H state observed in the absence of nucleotide or presence of non-hydrolyzable analogs (∼3% of the level observed in the presence of ATP) represents a mixture of inaccurate smFRET-Q state assignment and the formation of the locked BtuCD-F complex following transient release of B_12_ to bulk solution from BtuF* (*i.e.*, without movement into or through BtuCD). The trajectories of the transitions into the H state in the presence of WT BtuCD in the absence of nucleotide are predominantly consistent with formation of the locked complex, while those in the presence of AMPPNP are predominantly consistent with inaccurate mFRET-Q state assignment. This trend is consistent with our observations that the rate of the locking reaction is greatly reduced in the presence of AMPPNP34.

**Fig. S8.**
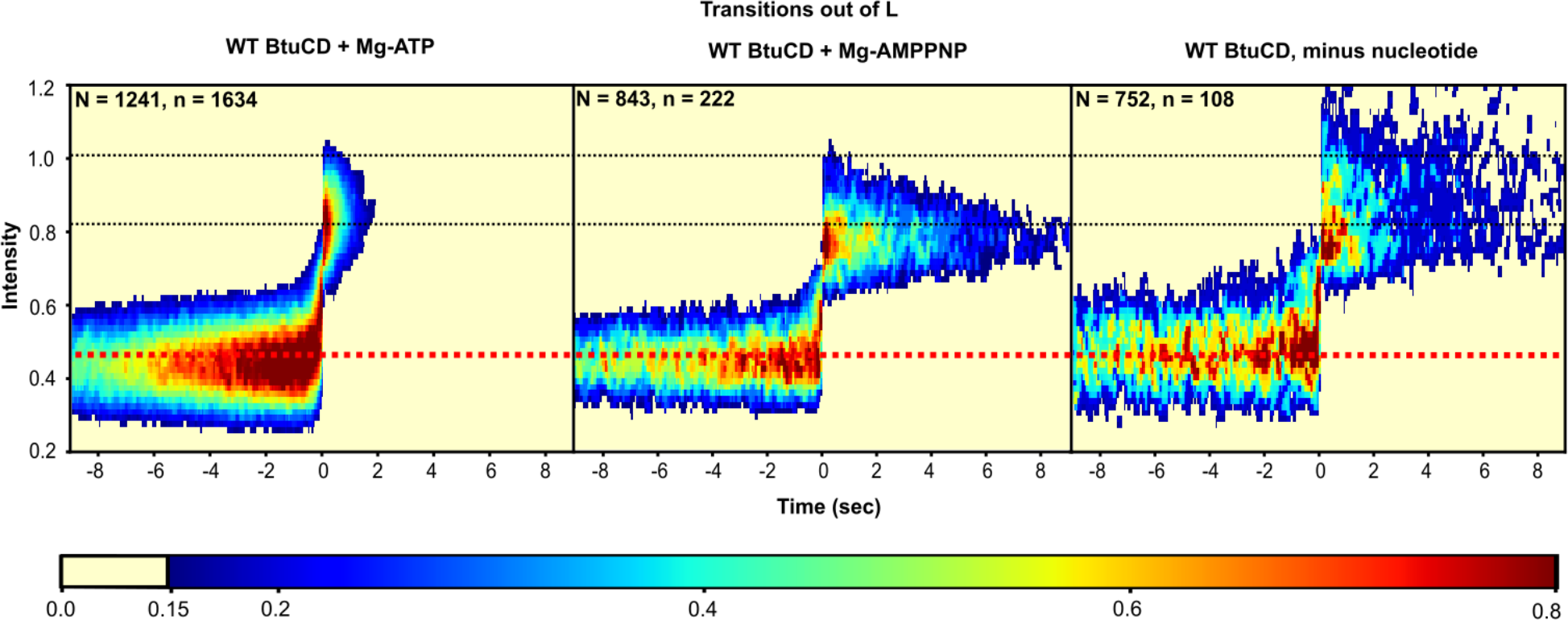
Synchronized plots for transitions out of the L state show evidence of substates with B_12_ at in intermediate positions. 2D contour plots showing complete trajectories equivalent to those shown in Fig. 2b in the main text. The systematically enhanced occupancy of smFRET-Q substates between the L and I states during the exit out of L suggest progressive movement of B_12_ from BtuF* to the internal cavity in BtuCD on a time-scale on the order of the 16.6 msec frame time.

**Fig. S9.**
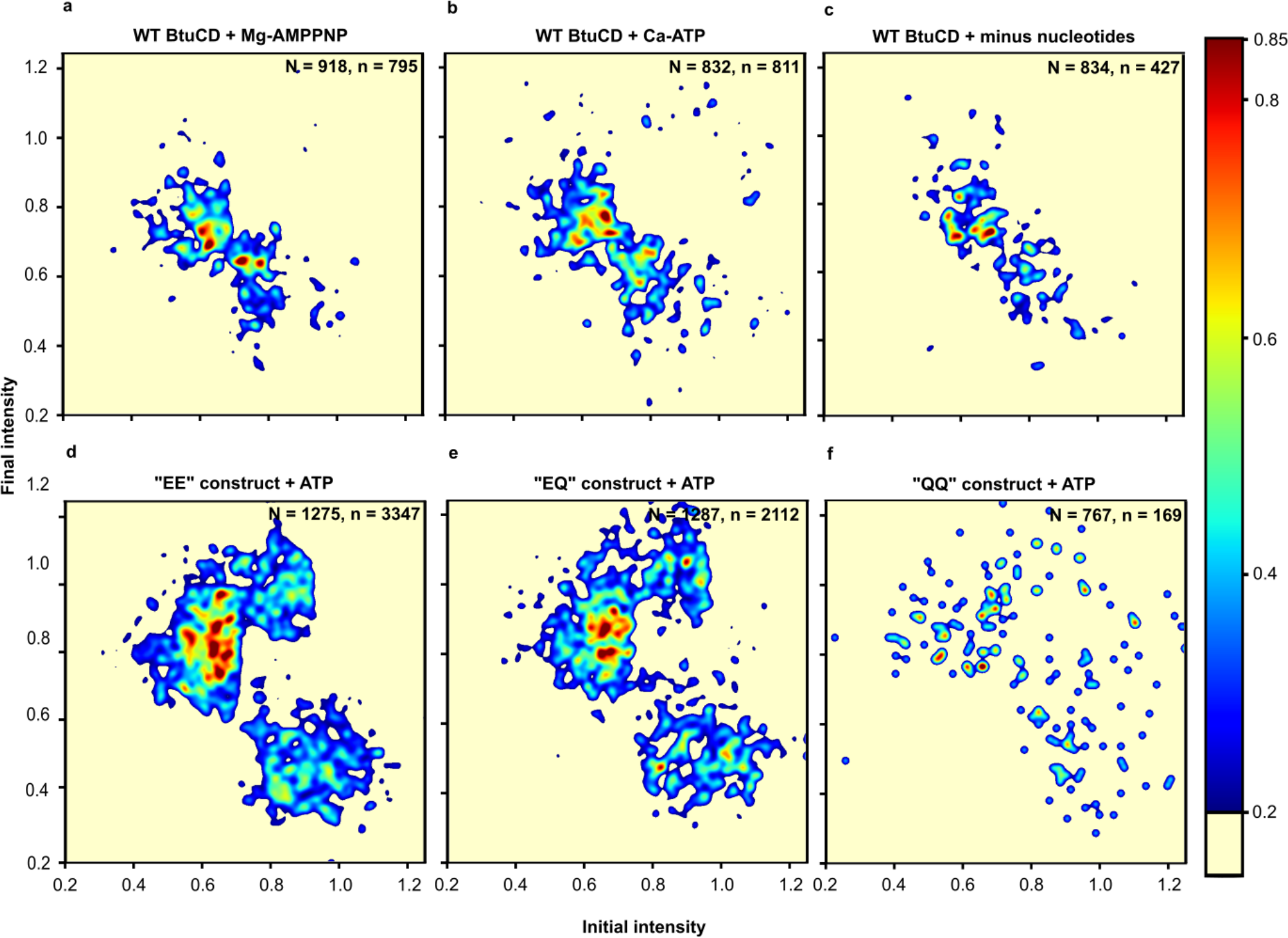
ATP hydrolysis drives unidirectional conformational transitions. Transition density plots equivalent to those shown in Fig. 2c but for experiments conducted in the presence of non-hydrolyzable analogs (**a-b**), in the absence of nucleotide (**c**), or with WT BtuCD replaced with tandem constructs (EE, EQ and QQ) (**d-f**). The plot in panel **e** is identical to the right plot in Fig. 2c but shown here again to facilitate comparison.

**Fig. S10.**
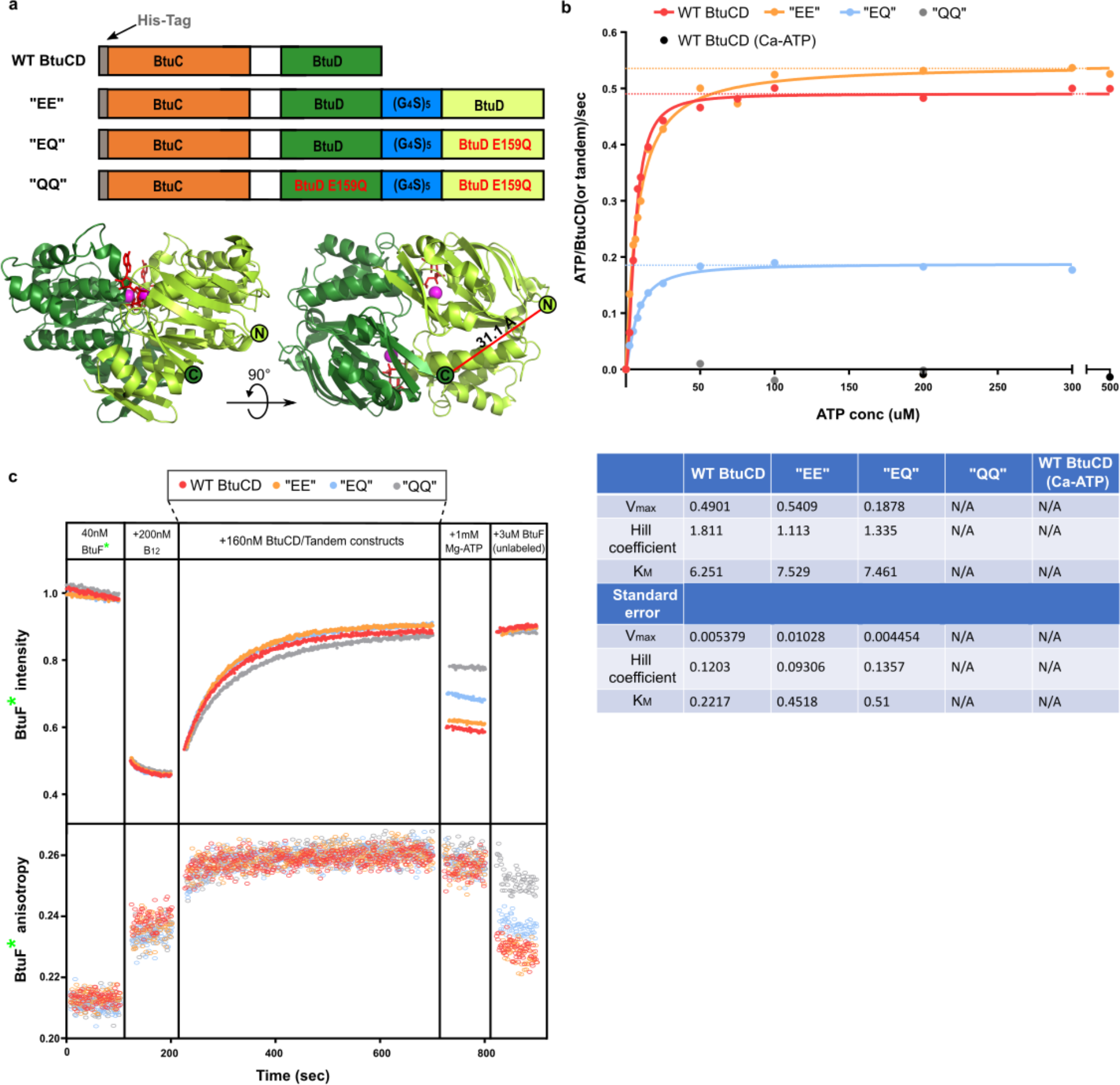
Covalently linked tandem BtuD_2_ constructs. **a,** DNA sequences comparing WT BtuCD construct and tandem constructs (upper panel), and ribbon diagrams of BtuD homodimer taken from BtuCD (PDB: 4FI3) (lower panel). The C-terminus of one monomer and N-terminus of the other monomer are shown. Inter-termini distance is 31.1Å. **b,** Initial ATP hydrolysis rate measured using the malachite green assay in the presence of 50 nM BtuCD (or tandem constructs), 1 µM BtuF, 2 µM B_12_, 2 mM Mg^2+^ (or Ca^2+^) and varied concentrations of ATP. Curve-fitting was done with the Michaelis–Menten equation. The results are shown in the lower panel. **c,** Ensemble experiments comparing WT BtuCD and tandem constructs. All constructs show equivalent binding behavior with BtuF* before the addition of ATP.

**Fig. S11.**
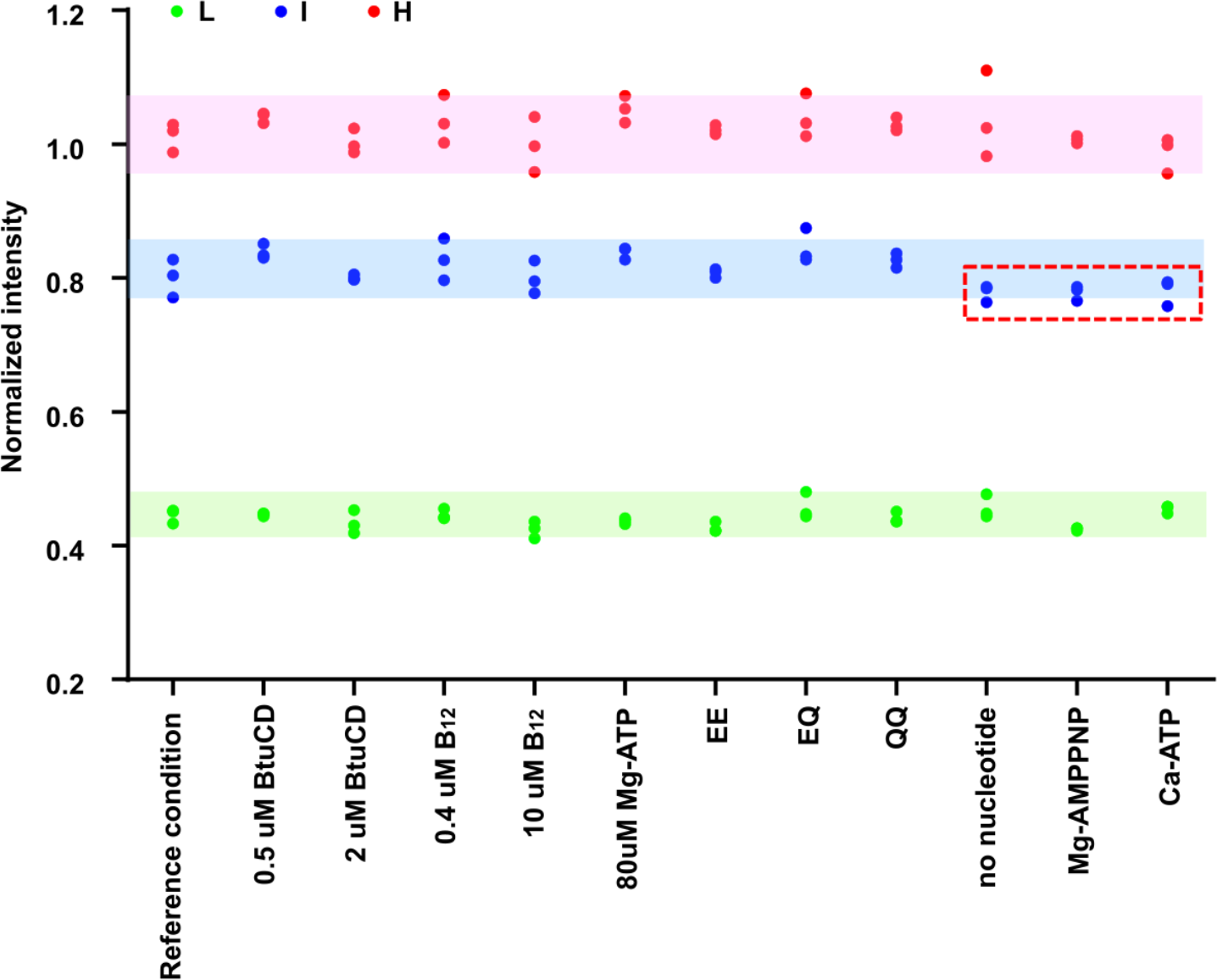
smFRET-Q intensity state means from HMM analysis. Triplicate measurements for all experimental conditions listed in **Supplementary Information Table 1** were independently analyzed using HMMs (see Methods). The derived means of the assigned L, I and H states are shown in green, blue and red. Experiments conducted with “no nucleotide”, “Mg-ATP” and “Ca-ATP” systemically show lower intensities for the I states compared to other conditions, as discussed in the main text and highlighted in red dash line box.

**Supplementary Information Table. 1.** smFRET experimental design. The table shows conditions for smFRET experiments that are discussed in the text and in the specified figures. Conditions that are different from the “reference condition”: 1 µM WT BtuCD, 1 µM B_12_ and 0.5 mM Mg-ATP are labeled in red.

| Experimental conditions | BtuCD (or tandem) | B<sub>12</sub> | Nucleotides | Figures |
| --- | --- | --- | --- | --- |
| Reference condition | 1 μM | 1 μM | 0.5 mM Mg-ATP | Figs. 1e-f, 2-3. Extended Data Fig. 8 |
| Varied BtuCD conc. | 0.5 μM | 1 μM | 0.5 mM Mg-ATP | Fig. 3a |
| | 2 μM | 1 μM | 0.5 mM Mg-ATP | |
| Varied B<sub>12</sub> conc. | 1 μM | 0.4 μM | 0.5 mM Mg-ATP | Fig. 3b |
| | 1 μM | 10 μM | 0.5 mM Mg-ATP | |
| Varied nucleotides | 1 μM | 1 μM | No nucleotide | Figs. 1e, g-h, 3c. Extended Data Figs. 7-9 |
| | 1 μM | 1 μM | 0.5 mM Mg-AMPPNP | |
| | 1 μM | 1 μM | 0.5 mM Ca-ATP | |
| | 1 μM | 1 μM | 80 μM Mg-ATP | |
| Tandem repeat constructs | 1 μM "EE" | 1 μM | 0.5 mM Mg-ATP | Figs. 3d Extended Data Figs. 7&9 |
| | 1 μM "EQ" | 1 μM | 0.5 mM Mg-ATP | |
| | 1 μM "QQ" | 1 μM | 0.5 mM Mg-ATP | |

